# Ripening, bursting, and synchronization of biomolecular condensates in a heterogeneous elastic medium

**DOI:** 10.1101/2023.05.27.542561

**Authors:** Lingyu Meng, Sheng Mao, Jie Lin

## Abstract

Biomolecular condensates play a crucial role in regulating gene expression, but their behavior in chromatin remains poorly understood. Classical theories of phase separation are limited to thermal equilibrium, and traditional methods can only simulate a limited number of condensates. In this paper, we introduce a novel mean-field-like method that allows us to simulate millions of condensates in a heterogeneous elastic medium to model the dynamics of transcriptional condensates in chromatin. Using this method, we unveil an elastic ripening process in which the average condensate radius exhibits a unique temporal scaling, ⟨*R*⟩ ∼ *t*^1*/*5^, different from the classical Ostwald ripening, and we theoretically derive the exponent based on energy conservation and scale invariance. We also introduce active dissolution to model the degradation of transcriptional condensates upon RNA accumulation. Surprisingly, three different kinetics of condensate growth emerge, corresponding to constitutively expressed, transcriptional-bursting, and silenced genes. Notably, multiple distributions of transcriptional-bursting kinetics from simulations, e.g., the burst frequency, agree with transcriptome-wide experimental data. Furthermore, the timing of growth initiation can be synchronized among bursting condensates, with power-law scaling between the synchronization period and dissolution rate. Our results shed light on the complex interplay between biomolecular condensates and the elastic medium, with important implications for gene expression regulation.

## I. INTRODUCTION

Biomolecular condensates are membraneless cellular compartments with many crucial physiological functions, e.g., stress adaptation, accelerating biochemical reactions, reducing noise [1–6]. Specifically, transcription-related condensates comprised of RNA polymerases (RNAPs) are observed in both prokaryotes and eukaryotes [7–12], suggesting an essential role of RNAP condensates in gene expression regulation. Biomolecular condensates are often liquid droplets forming via liquidliquid phase separation (LLPS) [2, 13], supported by their fluid-like behaviors [1, 14–16]. Classical LLPS theories focus on liquid droplets in a liquid environment at or evolving towards thermal equilibrium [13, 17, 18]. According to classical LLPS theories, molecules flow from small to big condensates to reduce the total surface energy, called Ostwald ripening. The outcome is a single large condensate because this configuration minimizes the surface energy. However, in many cases, the surrounding environments of biomolecular condensates are not simple viscous liquids, e.g., the nucleoplasm is filled with chromatin, and the cytoplasm contains cytoskeleton. It has been found that the elastic medium can interact with condensates and affect their growth and coarsening [19–26]. For example, light-activated condensates *in vivo* were observed to be constrained by the chromatin as the condensates could only stay in the chromatin-sparse region [23]. Soft matter experiments also found that the droplet-forming molecules flow from droplets in stiff medium to droplets in soft medium, suggesting a new driving force of coarsening due to elasticity beyond the classical LLPS theories [27, 28]. It is still unclear whether the elastic driving force generates any novel universality class of the ripening process beyond the Ostwald ripening.

Furthermore, numerous out-of-equilibrium processes consume energy inside living cells. In particular, the effects of active chemical reactions on the formation and morphologies of condensates have been intensely studied recently [29–31]. One notable function of active chemical reactions is to generate multiple stable coexisting condensates beyond the classical Ostwald ripening [1, 32– 35]. While active chemical reactions consume energies, such as ATP, alternative mechanisms to generate coexisting condensates have been proposed, e.g., through the nonlinear elasticity of surrounding network [20, 22, 25], which do not consume energy. In the meantime, other out-of-equilibrium processes also play crucial roles in condensate formation and dissolution. For example, recent experiments found that RNAP condensates can dissolve due to the accumulation of transcribed mRNAs, which is also an active process consuming energy [36].

In this work, we study the out-of-equilibrium dynamics of biomolecular condensates in a heterogeneous elastic medium. Regarding the theoretical tools, we introduce a novel mean-field-like model, which allows us to simulate millions of condensates simultaneously and significantly exceeds the computation capacity of traditional phase-field simulations [13]. We first study the case of a neo-Hookean elastic medium in which each condensate is confined by a constant local elastic pressure that can be different among condensates. Surprisingly, we find a new dynamical scaling of the average condensate radius ⟨*R*⟩∼ *t*^1*/*5^ during the ripening phase induced by heterogeneous elasticity, which we denote as elastic ripening. The 1*/*5 exponent is beyond the 1*/*3 exponent in the classical Ostwald ripening, and we derive its value based on principles of energy conservation and scale invariance. We also introduce nonlinear elasticity beyond the neo-Hookean model in which the ripening of condensates can be suppressed, and multiple condensates can coexist. However, within this nonlinear model, the system quickly reaches an equilibrium state without any temporal changes.

To incorporate biological activity, we assume that condensates dissolve at a rate proportional to their volume inspired by experiments in which RNAs can dissolve transcription-related condensates [36]. As a support of the active dissolution model, the simulated distribution of condensate lifetimes is similar to that of RNA polymerase II (Pol II) condensates in experiments [9]. Furthermore, depending on the local stiffness around condensates, the condensates can grow immediately after dissolution, grow intermittently, or be suppressed entirely, which we propose to correspond to constitutively expressed, transcriptional-bursting, and silenced genes. As another support of our theories, multiple simulated distributions of the kinetics of transcriptional-bursting genes, including the burst frequency and the burst size, agree with those of transcriptome-wide experimental data [37].

Surprisingly, a subset of the intermittently growing condensates with similar local stiffness can be synchronized so that they start growing simultaneously. We investigate the fraction of synchronized condensates and find a power-law scaling between the synchronization period with the dissolution rate. Our work reveals a new universality class for the ripening process of condensates in a heterogeneous elastic medium and uncovers the potential roles of chromatin elasticity in gene expression regulation.

### II. THE CONDENSATE GROWTH MODEL

We propose a mean-field-like model to describe the growth dynamics of condensates in a heterogeneous elastic medium (Figure 1). We assume that the condensate-forming biomolecule has a fixed concentration *c*_in_ inside the condensates and all condensates share a common outside concentration *c*_out_. This assumption is justified by phase-field simulations of condensate formation in which a virtually uniform outside concentration is observed (Supplementary Material and Figure S3). A uniform outside concentration is biologically reasonable because the typical protein diffusion constant *in vivo* is about 1 *μ*m^2^/s, which means that it takes seconds for a protein to fully explore the space inside a cell such as nucleus [38]. We assume near-equilibrium dynamics so that the changing rate of a condensate’s volume is proportional to the derivative of free energy with its volume,

**FIG. 1.**
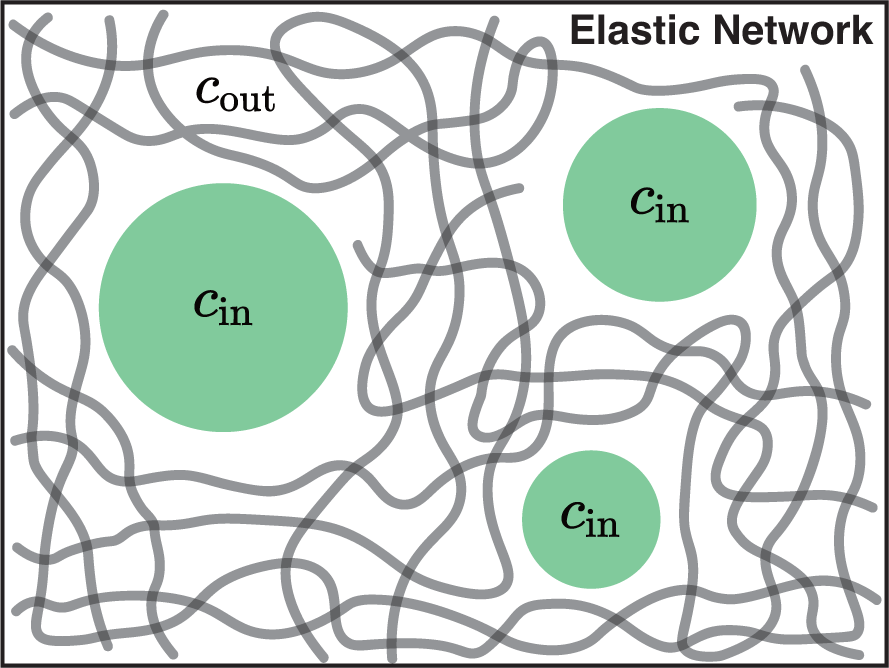
Biomolecular condensates grow in the interspace of a heterogeneous elastic medium. Each condensate has a fixed inside concentration *c*_in_ and a shared outside concentration *c*_out_. Condensates’ growth is suppressed by the surrounding elastic medium through a confining pressure, which can be different among condensates.

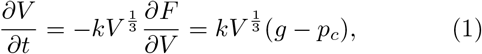

where *g* is the reduced free energy of condensate formation per unit volume, which we call condensing affinity, and

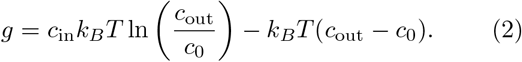

Here, *c*_0_ is the saturated concentration of phase separation and *g* = 0 when *c*_out_ = *c*_0_ as expected (see detailed derivations in Appendix A). The confining pressure *p*_*c*_ = *p*_*s*_ + *p*_el_. Here, *p*_*s*_ = 2*γ*(4*π/*3*V*)^1*/*3^ is the Laplace pressure and *γ* is the surface tension. *p*_el_ is the elastic pressure due to the surrounding medium, and we will explain its value later. The *V* ^1*/*3^ factor on the right side of Eq. (1) comes from the spherical geometry. The particle flux entering an absorbing sphere is proportional to its radius in three dimensions: *J*∝ 1*/R×R*^2^∼ *R* where 1*/R* is for the concentration gradient and *R*^2^ is for the surface area. As we show later, this *V* ^1*/*3^ factor is critical to obtain the correct scaling of the Ostwald ripening without heterogeneous elastic medium [13, 17, 18]. In the following, we non-dimensionalize the condensate growth model so that all the variables become dimensionless by selecting the energy unit *ϵ*_0_ = *k*_*B*_*T*, the length unit 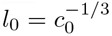 and the time unit 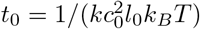.

>In this work, we mainly focus on the nucleation regime of phase separation because the average concentrations of condensate-forming molecules *in vivo* are typically far from the spinodal regime [39, 40]. We introduce a fixed number of nucleation sites with the nucleation radius *R*_*n*_, which can be related to the lengths of some specific DNA sequences, such as the promoters [12]. A condensate is initiated at a nucleation site if the condensing affinity overcomes the local Laplace and elastic pressure. Therefore, the larger *R*_*n*_, the easier the condensate to initiate growth. In the following numerical simulations, we set *c*_in_ = 10, the initial outside concentration *c*_out, ini_ = 2 and *γ* = 0.1. *c*_out_ is calculated as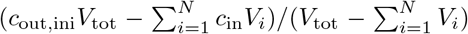 where *N* is the total number of nucleation sites. *V*_*i*_ is a condensate’s volume and *V*_tot_ is the total system volume, which we set as *V*_tot_ = 10^3^*N*. Neither *V*_*i*_ nor *V*_tot_ include the nucleation site volumes, which are typically very small. We define *R*_*c*_ ≡ 2*γ/g*_ini_, where *g*_ini_ is the initial condensing affinity so that *R*_*c*_ is the minimum nucleation radius for the condensate to grow at the beginning of the simulation in the absence of elastic pressure.

### III. ELASTIC RIPENING

In the following, we use the neo-Hookean elasticity to model the elastic medium in which the elastic energy cost is proportional to the condensate’s volume *F*_el_ = *EV* so the local elastic pressure *p*_el_ is *E* [25]. To mimic the heterogeneity of local stiffness in the nucleus, e.g., due to the spatial organization of euchromatin and heterochromatin, we assign each condensate a random *E*. In the case of Ostwald ripening, corresponding to the case of homogeneous *E*, small condensates shrink while big condensates grow as this ripening process reduces the overall surface energy. During the Ostwald ripening, the average and standard deviation of those survived condensates’ radii exhibit power scaling with time, ⟨*R*⟩∼ *σ*_*R*_∼ (*γt*)^1*/*3^ [13, 17, 18].

For the case of a heterogeneous elastic medium, we expect a similar ripening process in which condensates with large *E* shrink and condensates with small *E* grow since this can reduce the overall elastic energy. We simulate the condensate growth model and choose a uniform distribution of *E* from 0 to 2Ē where Ē is the average (Appendix B). Indeed, condensates initiated at nucleation sites with large *E* shrink during ripening (Figure S4), and nucleation sites with very large *E* may not initiate condensate growth throughout the simulations due to their strong elastic pressures.

Surprisingly, the system undergoing elastic ripening exhibits a novel power-law scaling between the average condensate radius and time, which crossovers to the Ostwald ripening at a later time (Figure 2a, b and Movie S1). Also, the standard deviation of the condensate radius exhibits a similar power-law scaling (Figure S5a). Moreover, the average and the standard deviation of the local elastic pressures also exhibit power-law scaling (Figure 2c and Figure S5b).To summarize,

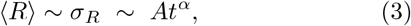

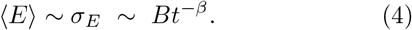

Here, 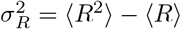 and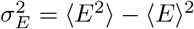. The average variable ⟨…⟩ is averaged over condensates weighted by their volumes, so the contribution of dissolved condensates is negligible. We also test the non-weighted average by excluding dissolved condensates explicitly and obtain similar results (Figure S6). *α* and *β* are the two power-law exponents. As we show later, *A* and *B* are the prefactors depending on the initial conditions.

**FIG. 2.**
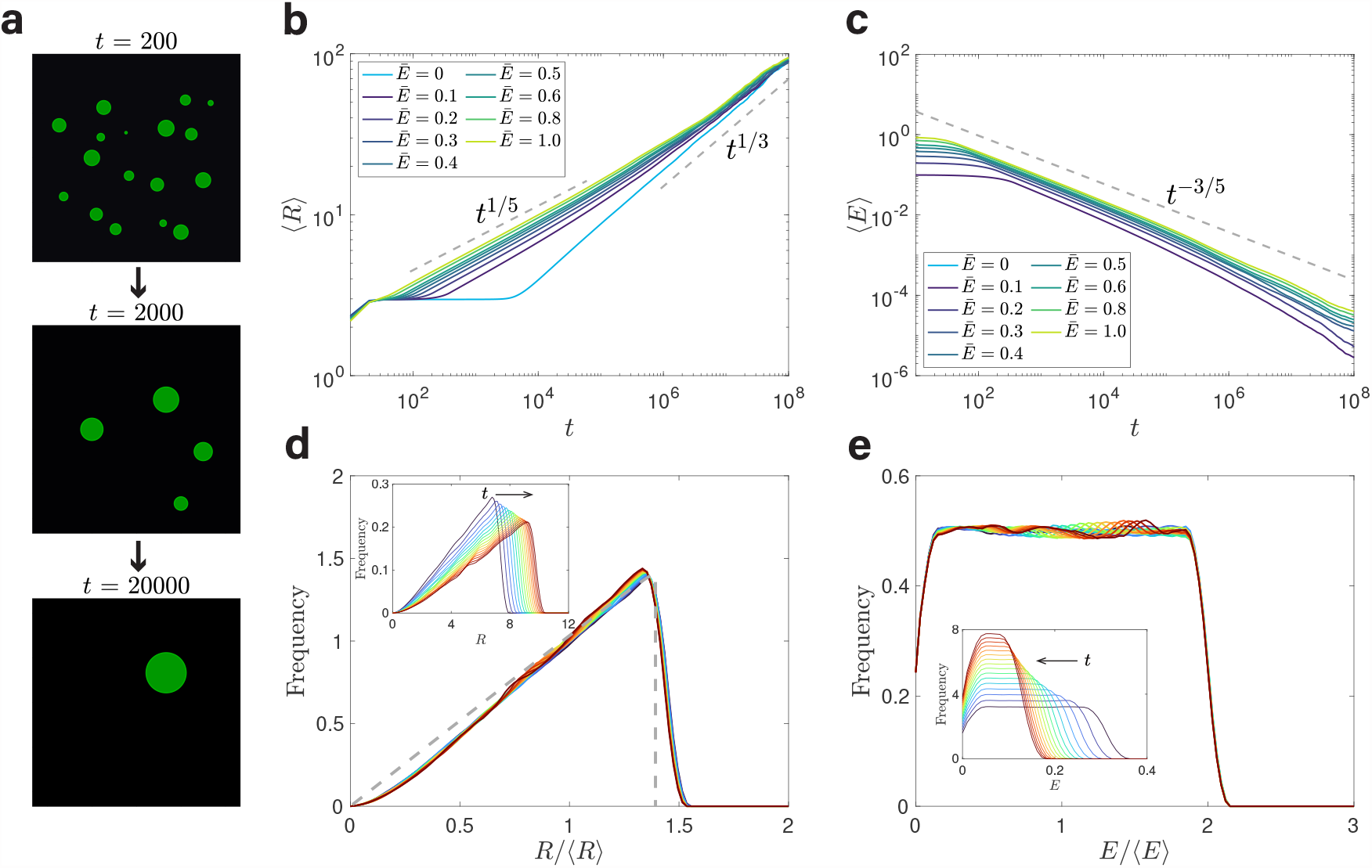
Elastic ripening in a heterogeneous neo-Hookean elastic medium. (a) Visualization of simulation results based on the condensate growth model. The systems undergo ripening until only one condensate survives. Note that the positions of these condensates are generated randomly and only a small subset of condensates are shown. (b) During the elastic ripening, the average radius exhibits a power-law scaling with time, ⟨*R* ⟩∼ *t*^1*/*5^. The elastic pressure *p*_el_ = *E* obeys a uniform random distribution in the range [0, 2Ē]At a later time, the elastic ripening crossovers to the Ostwald ripening in which ⟨*R*⟩∼ *t*^1*/*3^. (c) The average local elastic pressure ⟨*E*⟩ also exhibits a power law scaling with time, ⟨*E* ⟩∼*t*^*−*3*/*5^. (d) Distributions of the normalized radii *R/*⟨ *R*⟩ from *t* = 1000 to *t* = 4000 with a fixed interval converge to a universal distribution, exhibiting scale invariance. The gray dashed line is the theoretical prediction (Supplementary Material). Inset: the raw distributions of *R* at different times. (e) Distributions of the normalized radii *E/*⟨*E*⟩ from *t* = 1000 to *t* = 4000 with a fixed interval converge to a universal distribution, exhibiting scale invariance. Inset: the raw distributions of *E* at different times. In (a), (d), and (e), Ē= 1. In all figures, the radii of nucleation sites *R*_*n*_ = 1.5*R*_*c*_ where *R*_*c*_ = 2*γ/g*_ini_. For the case of Ē 0, we add randomness to *R*_*n*_ so that it is uniformly distributed from *R*_*c*_ to 2*R*_*c*_ to avoid deterministic dynamics. In (b) to (e), the total number of nucleation sites is 5 × 10^5^.

In the following, we theoretically derive the two exponents, based on the following relations:

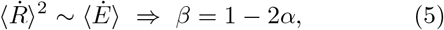

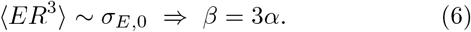

Eq. (5) comes from energy conservation: the dissipation power per unit volume 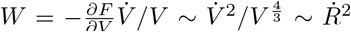 must be equal to the elastic energy changing rate per unit volume, which is *E*?. Eq. (6) comes from the scale invariance of the average elastic energy per condensate. *σ*_*E*m,0_ is the initial standard deviation of *E* with all nucleation sites contributing equally (excluding sites that never grow during the simulation). We remark that *σ*_*E*,0_ is the only scale of the initial *E* distribution (shifting the *E* distribution by a constant does not affect the dynamics of condensate growth). Based on Eqs. (5, 6), we obtain that *α* = 1*/*5 and *β* = 3*/*5. We can also obtain the expressions of the factor so that 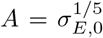 and 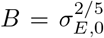 using Eqs. (5, 6). Therefore,

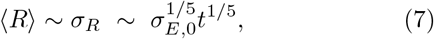

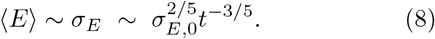

Our theoretical predictions are nicely confirmed for the average radius ⟨*R*⟩ (Figure 2b), the standard deviation of radius (Figure S5a), the average local elastic pressure *E* (Figure 2c), and the standard deviation of ⟨*E*⟩ (Figure S5b). We also verify the invariance of average condensate energy in Eq. (6) (Figure S7) and the expressions of the factors *A* and *B* (Figure S8). Our results do not depend on the distribution of *E* (Figure S9).

We have used the scale-invariance assumption in deriving Eq. (6), and the same power-law scaling of ⟨*R*⟩ and *σ*_*R*_ supports this idea. To explicitly test the scale-invariance assumption, we plot the distributions of the normalized condensates’ radii *R/* ⟨*R*⟩ at different times for the surviving condensates, and they indeed overlap, which means that the only length scale is the average radius ⟨*R*⟩ (Figure 2d). Notably, the distribution of *R/* ⟨*R*⟩ can be calculated semi-analytically, the gray dashed line in Figure 2d (Supplementary Material), although it depends on the distribution of *E* (Figure S10). Scale invariance is also observed for the distributions of *E/* ⟨*E*⟩ for the surviving condensates (Figure 2e). During the elastic ripening, the heterogeneity of local elastic pressures for the surviving condensates decreases while the heterogeneity of the radii for the surviving condensates increases. As a result, the Ostwald ripening eventually takes over the elastic ripening (Figure 2b).

## IV . BEYOND NEO-HOOKEAN

In a neo-Hookean medium, the elastic pressure is constant for each condensate, and the system undergoes ripening until only one condensate is left. It has been shown that a nonlinear elastic medium beyond neoHookean can suppress ripening [20, 22, 25]. To incorporate nonlinearity, we modify the elastic pressure so that it increases with the condensate radius *p*_el_ = *E* + *Y R* where *Y* is a constant, in agreement with recent molecular dynamical simulations [22]. We find that the average radius first follows the 1*/*5 elastic ripening scaling and then saturates to a plateau *R*_max_ (Figure 3 and Movie S2). From Eq. (1),<u>it is easy to</u> find *R*_max_ at equilibrium, 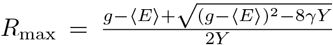. The system eventually reaches an equilibrium state where multiple condensates coexist with heterogeneous radii determined by the local elastic pressures.

**FIG. 3.**
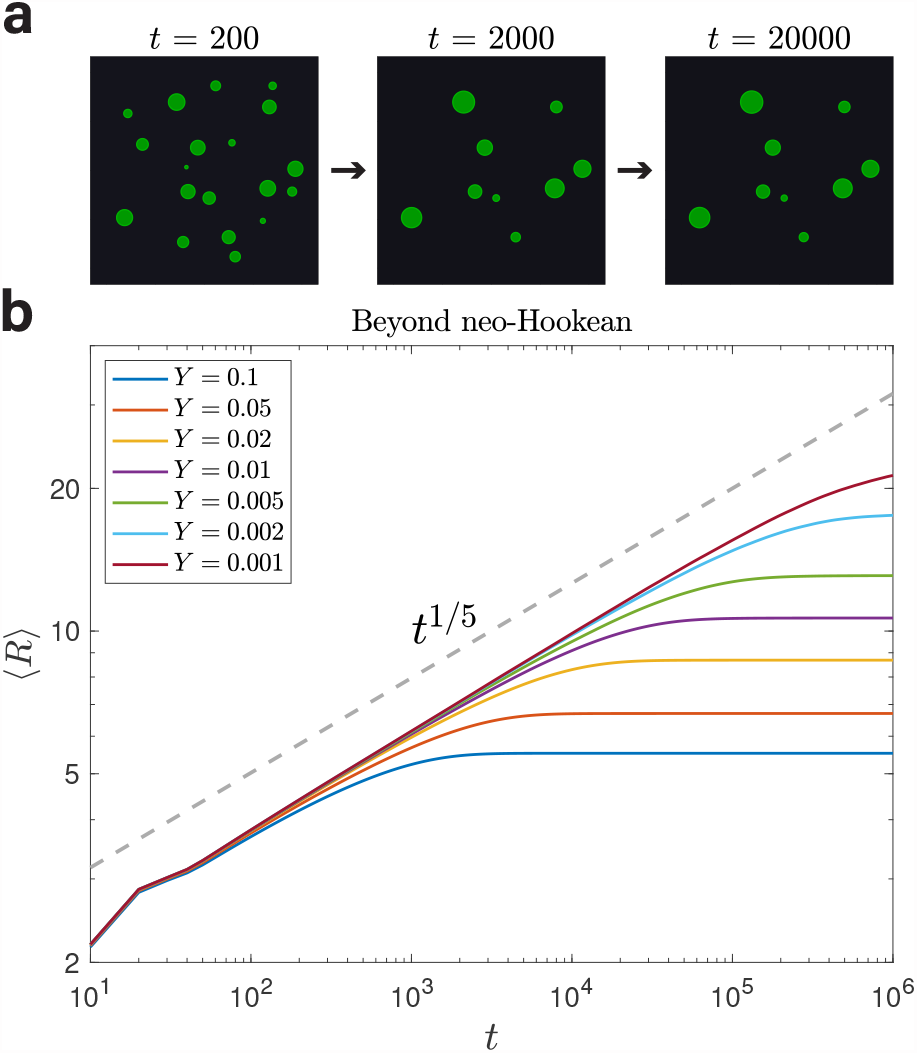
Elastic ripening is suppressed in an elastic medium beyond neo-Hookean elasticity. (a) Visualization of simulation results based on the condensate growth model. In this modified model, the elastic pressure increases with the condensate radius so that *p*_el_ = *E* +*Y R* where *E* obeys a uniform random distribution in the range [0, 2Ē] and *Y* is a constant. Here *Y* = 0.1. (b) The average radius ⟨*R*⟩ first exhibits the elastic ripening scaling and then reaches a plateau. In all panels, *E*Ē = 1 and the radii of nucleation sites *R*_*n*_ = 1.5*R*_*c*_. In (b), the total number of nucleation sites is 5 × 10^5^.

## V NONEQUILIBRIUM DYNAMICS OF ACTIVELY-DISSOLVING CONDENSATES

So far, we have analyzed the ripening dynamics of condensates as the system approaches thermal equilibrium, governed by the gradient of free energy. However, biological systems are often out of equilibrium due to energyconsuming processes. For example, transcriptional condensates can dissolve in the nucleus because of the accumulation of transcribed RNAs [36], and this active dissolution process makes the system depart from thermal equilibrium. To investigate the effects of active dissolution on the dynamics of transcriptional condensates, we introduce an active dissolution rate to each condensate so that they have a constant rate to dissolve per unit volume *k*_dis_, that is, the probability for condensate to dissolve within a short time window d*t* is *k*_dis_*V* d*t*. We assume the dissolution to be instantaneous because of fast protein diffusion [38, 41].

We first study the case of a heterogeneous neo-Hookean medium. To verify whether the active dissolution model is biologically reasonable, we calculate the distribution of condensate lifetimes from simulations and find that it is similar to the experimental data of Pol II condensates in live mouse embryonic stem cells from Ref. [9], supporting the validity of our model assumption (Figure S11). Interestingly, we find that the growth dynamics of condensates can be categorized into three cases. Condensates at nucleation sites with small local elastic pressures *E* grow immediately after dissolution (Figure 4a); condensates at nucleation sites with intermediate *E* grow after a delay since the last dissolution (Figure 4b); condensates at nucleation sites with large *E* never grow (Figure 4c). Intriguingly, these three cases can be mapped to three different gene expression kinetics: constitutively expressed genes, transcriptional-bursting genes, and silenced genes (Figure 4d). Understanding the kinetics of transcription has been a central challenge in the study of stochastic gene expression. Particularly, a bursting gene switches between the active and inactive states stochastically and is transcribed only when in the active state. Multiple mechanisms for transcriptional bursting have been proposed, such as histone modifications and transcription factor availability [37, 42–44]. Our results suggest that the local stiffness of chromatin provides a mechanical way to regulate transcription kinetics.

**FIG. 4.**
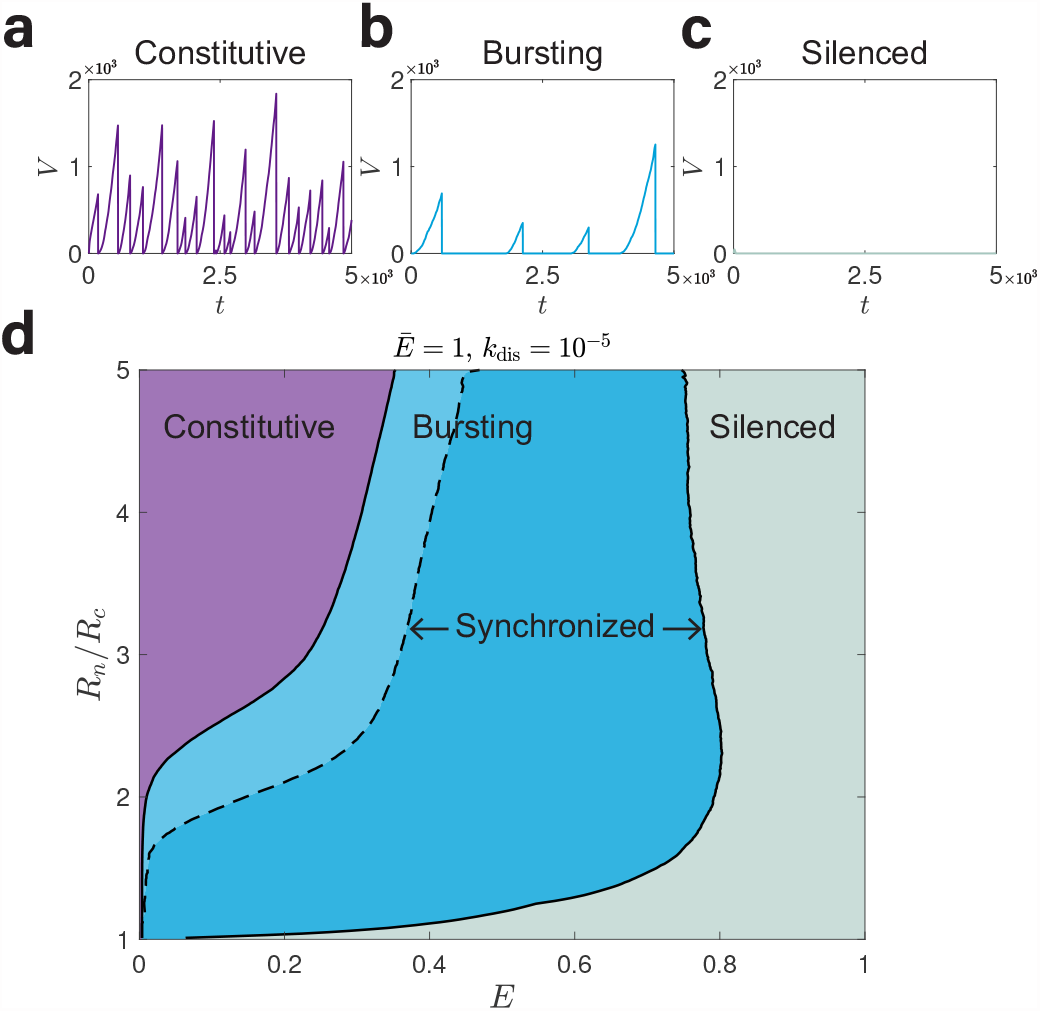
Dynamics of condensate growth and dissolution and its mapping to transcriptional kinetics. (a) Condensates with weak local stiffness, i.e., small *E*, grow immediately after dissolution and can be associated with constitutively expressed genes. (b) Condensates with intermediate local stiffness grow intermittently with a delay since the last dissolution and can be associated with transcriptional-bursting genes. (c) Condensates with strong local stiffness never grow, representing silenced genes that are not expressed. In (a) to (c), *R*_*n*_ = 8*R*_*c*_. (d) Phase diagram of condensate growth dynamics as a func-tion of *R*_*n*_ normalized by *R*_*c*_ and *E*. The purple, blue, and gray regions represent condensates associated with constitutively expressed, transcriptional-bursting, and silenced genes. The dark blue area represents the synchronized bursting con-densates. In all panels, Ē=1 *k*_dis_ = 10^*−*5^ and the number of nucleation sites *N* = 2000.

To test whether the transcriptional-bursting mechanism based on chromatin stiffness captures the main features of the bursting kinetics in experimental data, we compare the statistical properties of bursting condensates in our simulations to the transcriptome-wide allele-resolution experimental data in primary mouse fibroblasts from Ref. [37]. Four variables quantify the kinetics of a transcriptional-bursting gene: (1) the burst frequency, which quantifies how often the gene transitions to the active state and gets transcribed; (2) the burst size, which is the number of mRNAs produced during each burst; (3) the time until burst, which is the time interval between the finish of the last burst to the start of the next burst; (4) the time during burst, which is the time interval of a burst. Calculation details of bursting kinetics are included in Appendix B. Notably, the distributions of burst frequency, size, time until burst, and time during burst all qualitatively match the experimental data, particularly the skewness (Figure 5). These results suggest that our condensate growth model with active dissolution in a heterogeneous elastic medium captures some basic features of transcriptional kinetics *in vivo* (if not all). We also test other parameters for the simulations and obtain similar results (Figure S12).

**FIG. 5.**
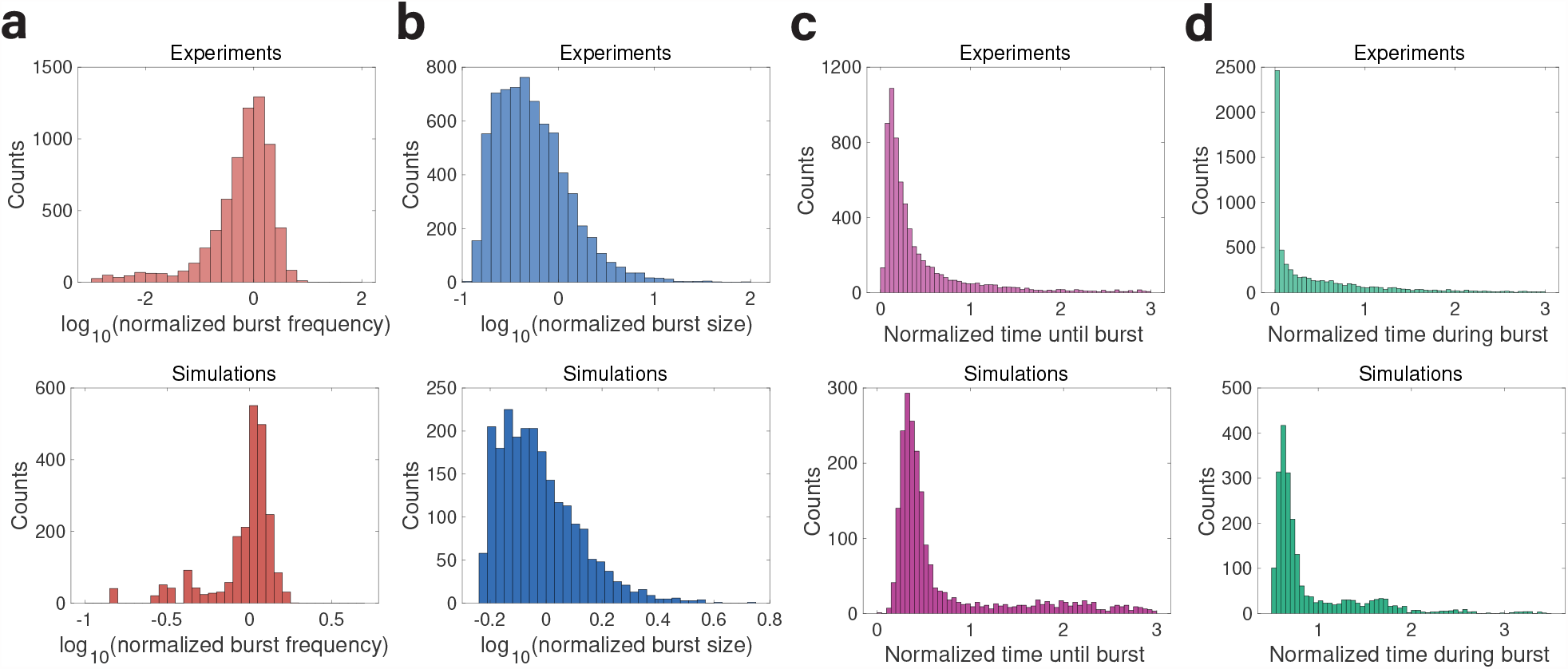
Comparison between experiments and simulations of transcriptional-bursting dynamics. The experimental data are transcriptome-wide distributions of transcriptional-bursting properties in primary mouse fibroblasts from Ref. [37]. Correspondingly, the simulated distributions are for the bursting condensates. (a) The count distribution of burst frequency from simulations of the condensate growth model (bottom) is similar to that from experiments (top). (b) Similar to (a) but for the distribution of burst size. (c) Similar to (a) but for the distribution of time until burst. (d) Similar to (a) but for the time distribution during a burst. In all panels, the distributions are normalized with their averages over all genes or condensates. In all panels of simulations, Ē=1, the radii of nucleation sites *R*_*n*_ = 5*R*_*c*_, *k*_dis_ = 10^*−*6^ and the number of nucleation sites *N* = 10^5^. Calculation details of bursting kinetics are included in Appendix B.

Furthermore, we observe a collective behavior of condensate growth: the initiation timings of a subset of bursting condensates are synchronized. When the nucleation radius *R*_*n*_ is small, the complete set of bursting condensates are synchronized (Figure 4d, Figure 6a, and Movie S3). As *R*_*n*_ increases, a subset of the bursting condensates remain synchronized. Eventually, the initiation timings of all condensates become completely uncorrelated (Figure 6b, c and Movie S4). Intuitively, when the nucleation radius *R*_*n*_ is small, the condensate has to overcome a large Laplace pressure to initiate growth. Therefore, condensates must wait until other condensates dissolve so that the outside concentration *c*_out_ is high enough to overcome the Laplace pressure. While for large *R*_*n*_, condensates can grow without the help of other condensates. We confirm that this phenomenon is generated by surface tension because the synchronized growth disappears in an artificial system without surface tension (Figure S13). This emerging synchronized dynamics of condensate growth suggests a mechanism of synchronized gene expression based on LLPS.

**FIG. 6.**
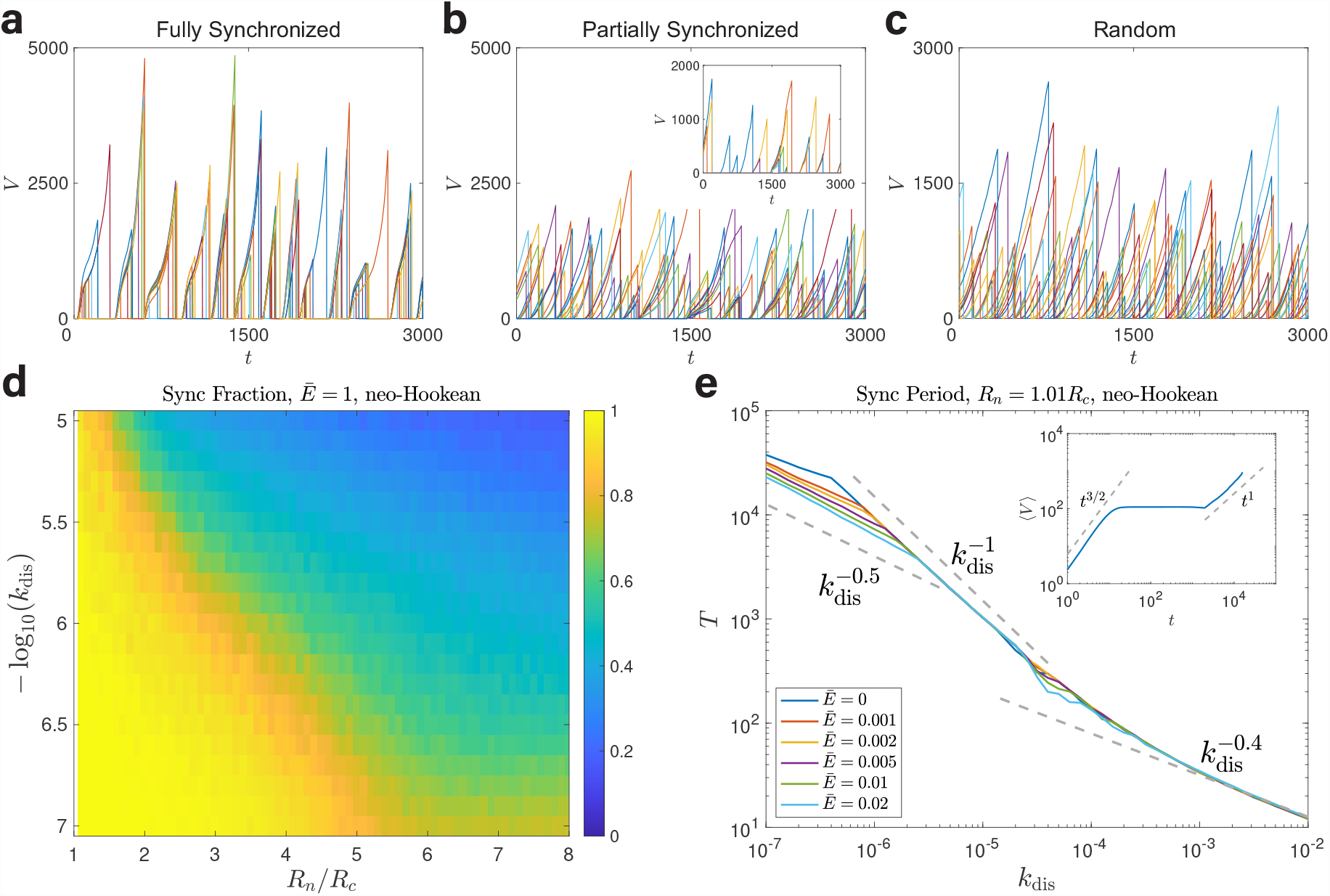
Synchronized growth of condensates in a neo-Hookean medium. (a) The temporal trajectories of condensate volumes *V* under a small *R*_*n*_, in which the initiation timings of all condensates are fully synchronized (excluding condensates that never grow during the simulation). Here, *R*_*n*_ = 1.2*R*_*c*_. (b) Here, *R*_*n*_ = 4*R*_*c*_. In this case, a subset of the bursting condensates are synchronized (inset). (c) Here, *R*_*n*_ = 10*R*_*c*_. In this case, the initiation timings of all condensates are completely uncorrelated. (d) The fraction of synchronized condensates as a function of *R*_*n*_*/R*_*c*_ and log_10_(*k*_dis_). (e) The period *T* exhibits piecewise power-law scaling with *k*_dis_. Inset: During one period of synchronization, the average volume over synchronized condensates ⟨*V* ⟩ ∼ *t* ^*3/2*^ during the growth phase, then reaches the plateau phase, and finally undergoes ripening with ⟨*V* ⟩ ∼ *t*. Here *k*_dis_ = 10^−7^ and Ē=0.01. In (e), *R*_*n*_ = 1.01*R*_*c*_ so that *f*_sync_ = 1 for the entire range of simulated *k*_dis_. In (a) to (d), Ē= 1. In all panels, the number of nucleation sites *N* = 2000.

We define an order parameter *f*_sync_ to represent the fraction of synchronized condensates excluding nucleation sites that never grow. When the fraction *f*_sync_ equals 1, all the condensates that can grow are synchronized. We obtain a heatmap of *f*_sync_ as a function of *R*_*n*_ and *k*_dis_ (Figure 6d). *f*_sync_ decreases gradually as *R*_*n*_ increases, indicating a continuous transition from fully synchronized to uncorrelated growth. Also, the *f*_sync_ decreases as the dissolution rate *k*_dis_ increases. This is because condensates release molecules to the environment more frequently when *k*_dis_ is large; therefore, the outside concentration *c*_out_ is high so that the coupling between condensates is weakened.

For the fully synchronized case (*f*_sync_ = 1), we predict that the period *T*, which is the time interval between two successive synchronized growth, should exhibit piecewise power-law scalings with the dissolution rate *k*_dis_. We estimate the period using *k*_dis_ ⟨*V*⟩ *T*∼ 1 where *V* is the average volume during one synchronization period in the limit of zero active dissolution rate. When the dissolution rate *k*_dis_ is large, the condensates dissolve during the initial growth phase. In this regime, ⟨*V* ⟩ ∼ *t*^3*/*2^ because *g p*_*c*_ is essentially constant [see Eq. (1) and the inset of Figure 6e]; therefore, 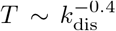. As *k*_dis_ decreases, the condensates can grow larger before they dissolve, and the average condensate size can reach the plateau phase before ripening in which ⟨*V*⟩ ∼ const; therefore, 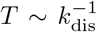. As *k*_dis_ further decreases, the system should undergo ripening. If the ripening is driven by elastic ripening, ⟨*V* ⟩ ∼ *t*^3*/*5^, and the period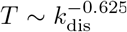. If the ripening is driven by Ostwald ripening, *V* ∼ *t*, and the period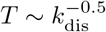.

To test our predictions, we simulate the condensate growth model by choosing *R*_*n*_ = 1.01*R*_*c*_ and a small Ē to ensure that *f*_sync_ = 1 in the entire range of simulated *k*_dis_. Our predictions are nicely confirmed (Figure 6e). We do not see a clear signature of the scaling 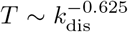 generated by elastic ripening in Figure 6e. We think that this is because the small range of *E* makes the regime of elastic ripening too short to observe (inset of Figure 6e). To test the existence of the scaling 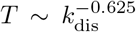, we increase the range of *E* and find a scaling behavior consistent with elastic ripening (Figure S14a). Our conclusions regarding the power-law scaling between *T* and the dissolution rate for the fully synchronized case are independent of the choice of *R*_*n*_ and *E*. Nevertheless, we note that the 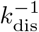 and 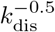 scalings occur only when *f*_sync_ = 1 (Figure S14b). This is because the plateau and ripening phases only apply to a closed system of synchronized condensates; that is, the outside concentration *c*_out_ is set by synchronized condensates only.

## VI. SYNCHRONIZED GROWTH BEYOND NEO-HOOKEAN

In the medium beyond neo-Hookean elasticity in which *p*_el_ = *E* + *Y R*, the phase space of bursting condensates shrinks while the phase space of constitutive condensates expands compared with the neo-Hookean model (Figure 7a, b). We calculate the synchronized fraction *f*_sync_ under different *Y* ‘s and find that *f*_sync_ decreases significantly when *Y* increases (Figure 7c). Because the increasing confining pressure limits the condensate growth, condensates tend to be smaller than the limiting case *Y* = 0, which means a higher outside concentration *c*_out_. Therefore, it is easier for condensate to initiate growth without waiting for other condensates’ dissolution. The synchronization period *T* also exhibits power-law scalings with the dissolution rate in the fully synchronized regime (Figure 7d). The main difference compared with the neoHookean model is that the 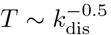 scaling disappears since ripening is suppressed by nonlinear elasticity. Further, the 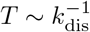 scaling does not require the condition *f*_sync_ = 1 anymore since the condensate volume will reach a plateau value in any case due to the increasing elastic pressure *p*_el_ = *E* + *Y R* (Figure 3 and Figure S15).

**FIG. 7.**
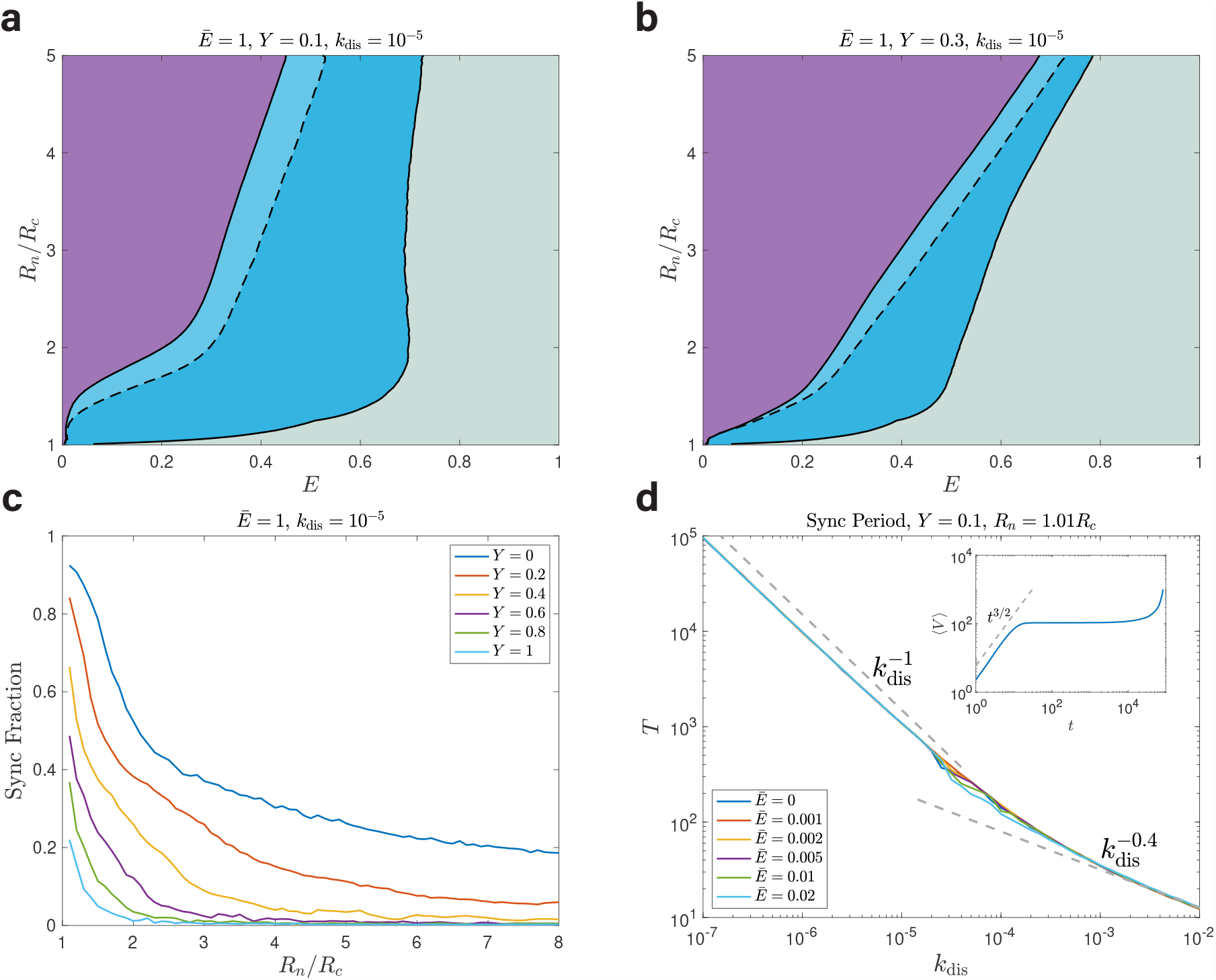
Synchronized growth of condensates in a medium beyond neo-Hookean. (a) The same phase diagram of condensate growth dynamics as Figure 4d but in a medium beyond neo-Hookean with *Y* = 0.1. (b) The same analysis as (a) with *Y* = 0.3. The fraction of synchronized condensates as a function of *R*_*n*_ given different *Y* ‘s. In (a) to (c), Ē=1, and *k*_dis_ = 10^*−*5^. The synchronization period *T* exhibits piecewise power-law scaling with *k*_dis_. Inset: during one synchronization period, the average volume over synchronized condensates ⟨*V*⟩∼ *t*^3*/*2^ during the growth phase, then reaches the plateau phase without the ripening phase. Here *k*_dis_ = 10^*−*7^ and Ē = 0.01. ⟨*V*⟩ increases at the end of the synchronization period because of the increasing outside concentration due to the early dissolution of some condensates, which nevertheless does not affect the scaling relation. In (d), *Y* = 0.1, *R*_*n*_ = 1.01*R*_*c*_ so that *f*_sync_ = 1 for the entire range of simulated *k*_dis_. In all panels, the number of nucleation sites *N* = 2000.

## VII. DISCUSSION

In this work, we have systematically investigated the dynamical behaviors of biomolecular condensates in a heterogeneous elastic medium. We introduce a meanfield-like model to investigate the condensate growth dynamics, allowing us to track millions of condensates simultaneously. Using this novel numerical method, we find a new dynamical scaling for the elastic ripening, ⟨*R*⟩∼ *t*^1*/*5^, and theoretically derive the origin of the 1*/*5 exponent based on energy conservation and scale invariance. Furthermore, the heterogeneity of local elastic pressure decreases over time and exhibits power-law scaling as well, *σ*_*E*_ ∼ *t*^−3*/*5^. Ripening is suppressed in an elastic medium beyond neo-Hookean elasticity, and multiple condensates can coexist at equilibrium. Our theoretical predictions and numerical simulations nicely agree with each other.

To incorporate biological activity, we also introduce a constant dissolution rate per unit volume to each condensate to model the dissolution of transcriptional condensates *in vivo* due to RNA accumulation [36]. This active dissolution process drives the system out of equilibrium. As evidence of the validity of the active dissolution model, the simulated distribution of condensate lifetimes is similar to experimental data [9]. Intriguingly, the temporal growth patterns of condensates resemble gene expression dynamics. Condensates in soft regions with weak local stiffness keep growing and continue to grow immediately after dissolution, corresponding to constitutively expressed genes. Meanwhile, condensates with stronger local stiffness initiate growth after a delay since the last dissolution, corresponding to transcriptional-bursting genes, which switch between active and inactive states and initiate transcription only in the active state. If the local stiffness is too strong, condensates can never grow, corresponding to silenced genes that are not expressed. Surprisingly, the simulated distributions of multiple characteristics of transcriptional burst, including the burst frequency and size, nicely reproduce the transcriptome-wide experimental distributions [37]. Our results suggest that the local mechanical properties of chromatin play a key role in regulating gene expression kinetics, which can be another layer of regulation on top of the compartmentalization of euchromatin and heterochromatin.

Notably, the timing of the growth initiation of bursting condensates can be synchronized. We remark that this is entirely a nonequilibrium effect due to the active dissolution process and finite surface tension. The fraction of synchronized condensates depends on the nucleation radius and dissolution rate. For the fully-synchronized cases, the period of synchronized growth exhibits piecewise power-law scalings with the dissolution rate. We theoretically derive the power-law exponents and show that it is related to the power-law scalings of average condensate sizes with time. In an elastic medium beyond neo-Hookean, synchronized growth is suppressed, and the results are qualitatively similar.

Some questions remain regarding the connection of our results to condensates in natural biological systems. In the experiments by Cho et al., though the lifetime distribution of transient condensates can be captured by our model, stable and large condensates that virtually do not dissolve within the experimental time window coexist with smaller condensates that dissolve frequently [9]. This indicates that the nonequilibrium dissolution process of biomolecular condensates can be more complex than our simplified assumptions. The coalescence of condensates is also not included in our model. Recently, Lee et al. found a new dynamical scaling of ripening generated by the coalescence of subdiffusive condensates [23]. In the future, it will be interesting to explore the interference of elastic ripening, driven by the gradient of local stiffness and the subdiffusion of condensates themselves.

## ACKNOWLEDGMENTS

We thank Fanlong Meng, Zhi Qi, and Yiyang Ye for useful discussions related to this work. The research was funded by National Key R&D Program of China (2021YFF1200500) and supported by grants from Peking-Tsinghua Center for Life Sciences.

L.M. conceived, designed, and carried out the theoretical and numerical part of this work. S.M. conceived and designed the theoretical part of this work. J.L. conceived, designed, and carried out the theoretical and numerical part of this work. All the authors contributed to the preparation of the manuscript.

## APPENDIX A DETAILED DERIVATION OF THE CONDENSATE GROWTH MODEL

We compute the condensing affinity *g* in the condensate growth model by considering a small change of the condensate volume *V* with its inside concentration fixed at *c*_in_:

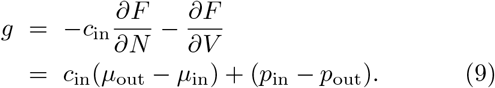

Here, *μ*_in_ and *μ*_out_ are the chemical potentials of the condensate-forming molecules inside and outside the condensate. *p*_in_ and *p*_out_ are the pressures inside and outside the condensate. Next, we use the Gibbs-Duhem equation, *N* d*μ* = −*S*d*T* + *V* d*p* [45]. Therefore, *μ*_in_ = *μ*_0_ + (*p*_in_ − *p*_0_)*/c*_in_ where *μ*_0_ is the chemical potential of the condensate-forming molecules and *p*_0_ is the pressure at equilibrium in the thermodynamic limit.

Regarding the chemical potential and the pressure outside condensates, we have d*μ*_out_ = *k*_*B*_*T* d*c*_out_*/c*_out_ and d*p*_out_ = d*μ*_out_*/c*_out_. Here, we have used the dilute solution assumption for d*μ*_out_ and the Gibbs-Duhem relation for d*p*_out_. Integrating *c*_out_ from *c*_0_, the saturation concentration at equilibrium in the thermodynamic limit, we obtain *μ*_out_ = *μ*_0_ + *k*_*B*_*T* ln(*c*_out_*/c*_0_) and *p*_out_ = *p*_0_ + *k*_*B*_*T* (*c*_out_ *c*_0_). Combining all the above results, we obtain

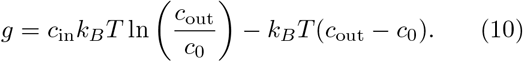

## APPENDIX B DETAILS OF NUMERICAL SIMULATIONS AND DATA ANALYSIS

We perform numerical simulations by solving Eq. (1) using the explicit Euler method on MATLAB. In Figure 4d, Figure 7a, and Figure 7b, a condensate is mapped to a constitutively expressed gene if more than half of its growths initiate within a dimensionless time 100 since the last dissolution. Meanwhile, a condensate is mapped to a silenced gene if it never grows during the simulation. Finally, the rest condensates correspond to transcriptionalbursting genes.

In Figure 5a, a condensate’s burst frequency is calculated as the inverse of the average interval between successive growth initiations. In Figure 5b, a condensate’s burst size is approximated as the product of its final volume before dissolution and half of its lifetime. In Figure 5c, the time until burst is calculated as the average interval from dissolution to the next initiation. In Figure 5d, the time during burst is calculated as its average lifetime. In all panels of Figure 5 for the simulations, we exclude the data of small condensates whose burst size is smaller than a threshold 10^6^.

In Figure 4d, Figure 6, and Figure 7, we label the condensate that can grow with the largest *E* as the first synchronized condensate. We then scan the condensates that can grow from the largest *E* to the smallest *E*, and the condensate is labeled as synchronized if over 80% of its initiation timings overlap with the growth timing of the latest synchronized condensate. After all the synchronized condensates are identified, we calculate the period by finding the average interval between two successive peaks of the distribution of initiation timings for all synchronized condensates.

In Figure 6e and 7d, in the case of Ē = 0, we add a small noise ±0.001*R*_*n*_ to the nucleation radius to avoid deterministic dynamics. In the inset of Figure 6e, we exclude the ⟨*V*⟩ data when the survived condensates’ number is smaller than 200 below which finite size effects are significant.

Movie S1: Condensates undergo elastic ripening in a heterogeneous neo-Hookean elastic medium until only one condensate exists. The simulation result is the same as Figure 2a in the main text. In the movie, we show a finite number of nucleation sites, randomly and uniformly chosen from the distribution of *E*. The same protocol applies to other movies.

Movie S2: Elastic ripening is suppressed in an elastic medium beyond neo-Hookean elasticity. The simulation result is the same as Figure 3a in the main text.

Movie S3: All actively-dissolving condensates are synchronized when *R*_*n*_ is small. Simulation parameters are the same as in Figure 5a in the main text.

Movie S4: All actively-dissolving condensates grow randomly when *R*_*n*_ is large. Simulation parameters are the same as in Figure 5c in the main text.

## Notes

### Competing Interest Statement

The authors have declared no competing interest.

### Summary of Updates

Figure 5 and relevant discussions added to compare our model to the experimental data. Supplemental files updated.

