## Supplementary Material for "Ripening, bursting, and synchronization of biomolecular condensates in a heterogeneous elastic medium"

Lingyu Meng,[1] Sheng Mao,[2] and Jie Lin[1,3]
[1] *Peking-Tsinghua Center for Life Sciences, Academy for Advanced Interdisciplinary Studies, Peking University, Beijing 100871, China*
[2] *Department of Mechanics and Engineering Science, BIC-ESAT, College of Engineering, Peking University, Beijing 100871, China*
[3] *Center for Quantitative Biology, Academy for Advanced Interdisciplinary Studies, Peking University, Beijing 100871, China*
(Dated: July 17, 2023)

### A. Details of phase-field simulation

We numerically simulate a two-fluid model similar to Model H involving diffusion and advection [1–4]. In the two-fluid model, the biomolecule density field $\phi$ is spatially dependent with its value between 0 and 1. The velocity field is $\mathbf{v}$. The dynamics of the biomolecule density and velocity field follows [3]

$$\frac{\partial \phi}{\partial t} = -\nabla \cdot (\phi \mathbf{v}) + \nabla\Big(\frac{\phi(1-\phi)^2}{\zeta}(\nabla \cdot \mathbf{\Pi})\Big), \tag{1}$$

$$-\nabla \cdot \mathbf{\Pi} - \nabla p + \eta \nabla^2 \mathbf{v} = 0. \tag{2}$$

Here, $\zeta$ is the friction constant between biomolecules and solvent, and $\eta$ is the viscosity. The pressure $p$ is determined by the incompressible condition: $\nabla \cdot \mathbf{v} = 0$. The osmotic stress tensor $\mathbf{\Pi}$ is determined by the biomolecule free energy $f(\phi)$, $\nabla \cdot \mathbf{\Pi} = \phi \nabla f'(\phi)$ where $f(\phi) = f_0(\phi) + \frac{C}{2}(\nabla \phi)^2$ and $C$ is a constant. We use the Flory-Huggins free energy density: $f_0(\phi) = \epsilon_0(\phi \ln(\phi) + (1-\phi)\ln(1-\phi) + \chi\phi(1-\phi))$ and take $\chi = 3$ in Figure S3. We non-dimensionalize our model with the unit of energy density as $\epsilon_0$, the time unit as $t_0 = \eta/\epsilon_0$ and the length unit as $l_0 = \sqrt{\eta/\zeta}$. In our simulation, we take $C/\epsilon_0 l_0^2 = 1$. We perform numerical simulations in a 2D $255 \times 255$ grid by solving the two-fluid model using the explicit Euler method with the periodic boundary condition on MATLAB. The grid size is $\Delta l = 0.25$. The time interval for the simulation is $\Delta t = 0.001$. The initial average density in Figure S3 is 0.45, and we add a $\pm 0.01$ noise to trigger phase separation.

### B. Derivation of the radius distribution for elastic ripening

The density $n$ of the condensate radius $R$ obeys the following continuous equation [5]:

$$\frac{\partial n}{\partial t} + \frac{\partial}{\partial R}(nv) = 0, \tag{3}$$

where $v$ is the changing rate of condensate radius, i.e., $\mathrm{d}R/\mathrm{d}t$. We denote the time-dependent average radius as $R_t$ and let $x = R/R_t$. Guided by the simulation results, we propose the following ansatz for the expression of $v$:

$$v = R_t^k h(x), \tag{4}$$

where $k$ is an exponent to be determined. Since $n$ is the density of radius per unit volume in three dimensions, we have

$$n = R_t^{-4} f(x). \tag{5}$$

Plugging Eqs. (4, 5) to Eq. (3), we obtain the following equations

$$R_t^{-k}\dot{R}_t = C, \tag{6}$$

$$C[xf'(x) + 4f(x)] = \frac{\partial}{\partial x}[f(x)h(x)] \tag{7}$$

From Eq. (6), we obtain the power-law scaling $R_t \sim t^{1/(-k+1)}$.

In our mean-field-like model with neo-Hookean elasticity, the changing rate of condensate radius is

$$v = \frac{\mathrm{d}R}{\mathrm{d}t} = \frac{1}{4\pi R}(g - E). \tag{8}$$

Here we have ignored the Laplace pressure since we focus on the elastic ripening. From the simulation results with uniform random $E$, we observe

$$\frac{E}{E_t} = e_{\max} - \left(\frac{R}{R_t}\right)^2 = e_{\max} - x^2, \tag{9}$$

where $E_t$ is the average local elastic pressure and $e_{\max}$ is the maximum $E/E_t$ (Figure S1). We note that this relation is not universal against the distribution of $E$.

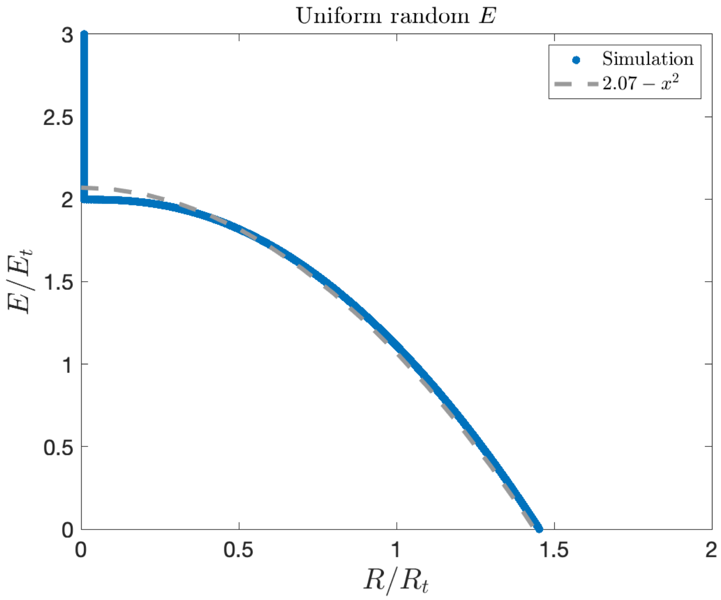


FIG. S1. Relation between $E/E_t$ and $R/R_t$ given a uniformly distributed $E$. The blue dots are simulation results and the gray dashed line is the fitting according to Eq. (9). In this figure, $\bar{E} = 1$.

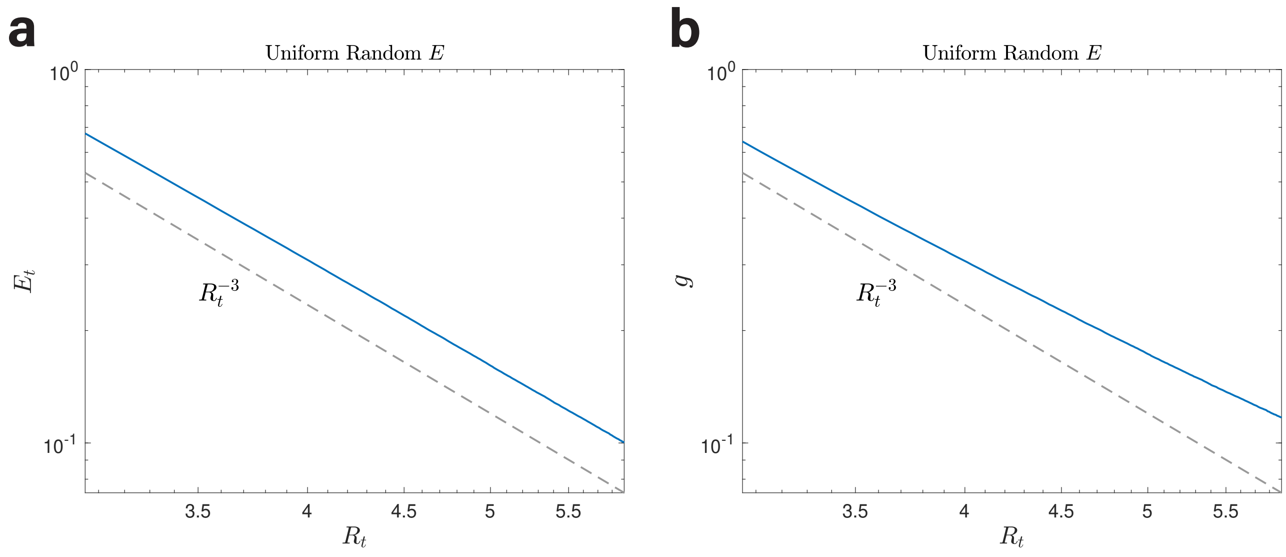


FIG. S2. $E_t$ and $g$ have power-law scaling with $R_t$. (a) $E_t$ is proportional to $R_t^{-3}$ in the numerical simulations, satisfying scale invariance. (b) $g$ is also approximately proportional to $R_t^{-3}$ in the numerical simulations, satisfying scale invariance. In this figure, $\bar{E} = 1$.

Next, we use the condition of scale invariance so that

$$E_t = \frac{C_1}{R_t^3}, \tag{10}$$

confirmed by the numerical simulations (Figure S2a). To achieve scale invariance, $g$ must have the same scaling as $E_t$:

$$g = \frac{C_2}{R_t^3}, \tag{11}$$

also confirmed by the simulation results (Figure S2b). Thus, we obtain the expression of $v$ as

$$v = \frac{1}{4\pi R} \frac{C_2 - C_1[e_{\max} - (R/R_t)^2]}{R_t^3} = R_t^{-4} h(x), \tag{12}$$

where

$$h(x) = \frac{1}{4\pi}\left(\frac{C_2 - e_{\max} C_1}{x} + C_1 x\right). \tag{13}$$

Eq. (12) suggests that $k = -4$, which means that $R_t \sim t^{1/5}$, in agreement with our conclusions in the main text. Finally, we obtain the distribution function of condensate radius $f(x)$ by solving Eq. (7):

$$f(x) \propto x \left[(e_{\max} C_1 - C_2) - (C_1 - 4\pi C)x^2\right]^{\dfrac{10\pi C - C_1}{C_1 - 4\pi C}}, \tag{14}$$

Compared with the simulation results, we find $e_{\max} = 2.07$, $C = 0.61$, $C_1 = 19.10$ and $C_2 = 17.36$. We plot the predicted distribution function $f(x)$, which agree reasonably well with the numerical results (Figure 2d in the main text).

From the simulation results, we find that $f(x)$ appears to be linear in a wide range of $x$. Intriguingly, we note that this suggests that $C$ and $C_1$ are not independent constants. Assuming a linear form of $f(x)$, we simplify Eq. (7) to

$$h(x) + x h'(x) = 5Cx, \tag{15}$$

with the solution

$$h(x) = \frac{A}{x} + \frac{5C}{2} x, \tag{16}$$

where $A$ is an arbitrary constant. The structure of Eq. (16) is similar to Eq. (13). By matching the linear term, we calculate $C = C_1/10\pi \approx 0.6081$ in Eq. (16) from the fitted $C_1$ in Eq. (13), which is very close to the fitted $C = 0.61$.

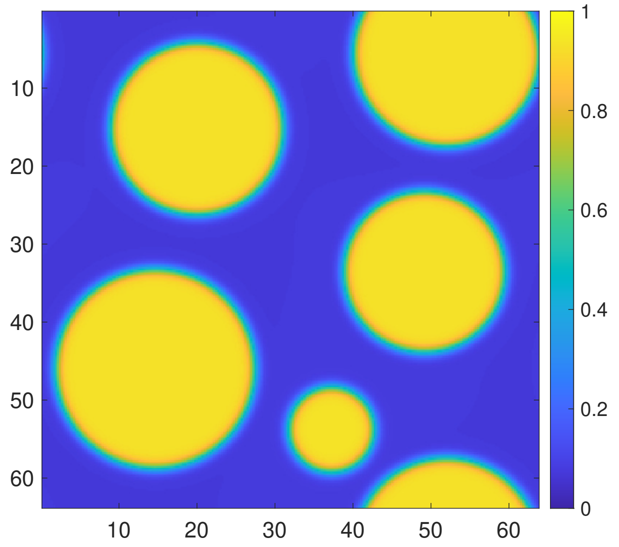


FIG. S3. Simulation of phase separation using phase-field model in which the outside concentration is uniform without obvious gradient.

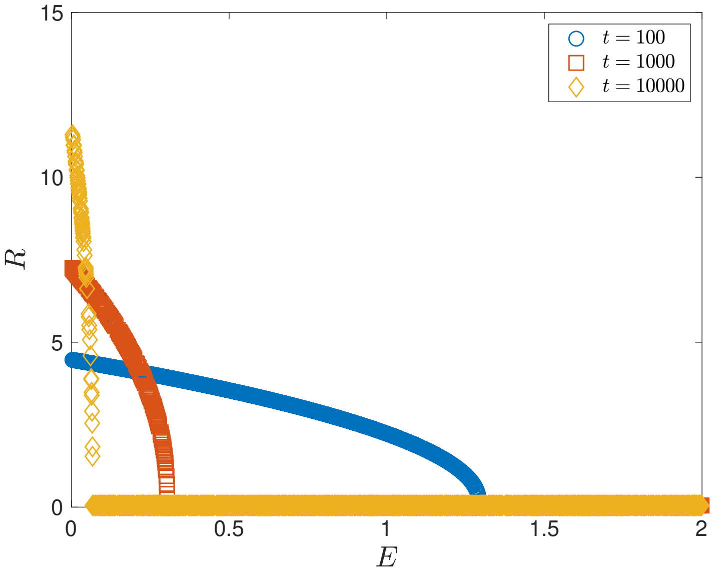


FIG. S4. The condensate radius $R$ v.s. the local elastic pressure $E$ during the elastic ripening at different times. Condensates with large $E$ shrink while condensates with small $E$ grow. In this figure, $E$ obey a uniform random distribution in the range $[0, 2\bar{E}]$ and $\bar{E} = 1$. $R_n = 1.5R_c$ with $R_c = 2\gamma/g_{\text{ini}}$. The number of nucleation sites $N = 2000$.

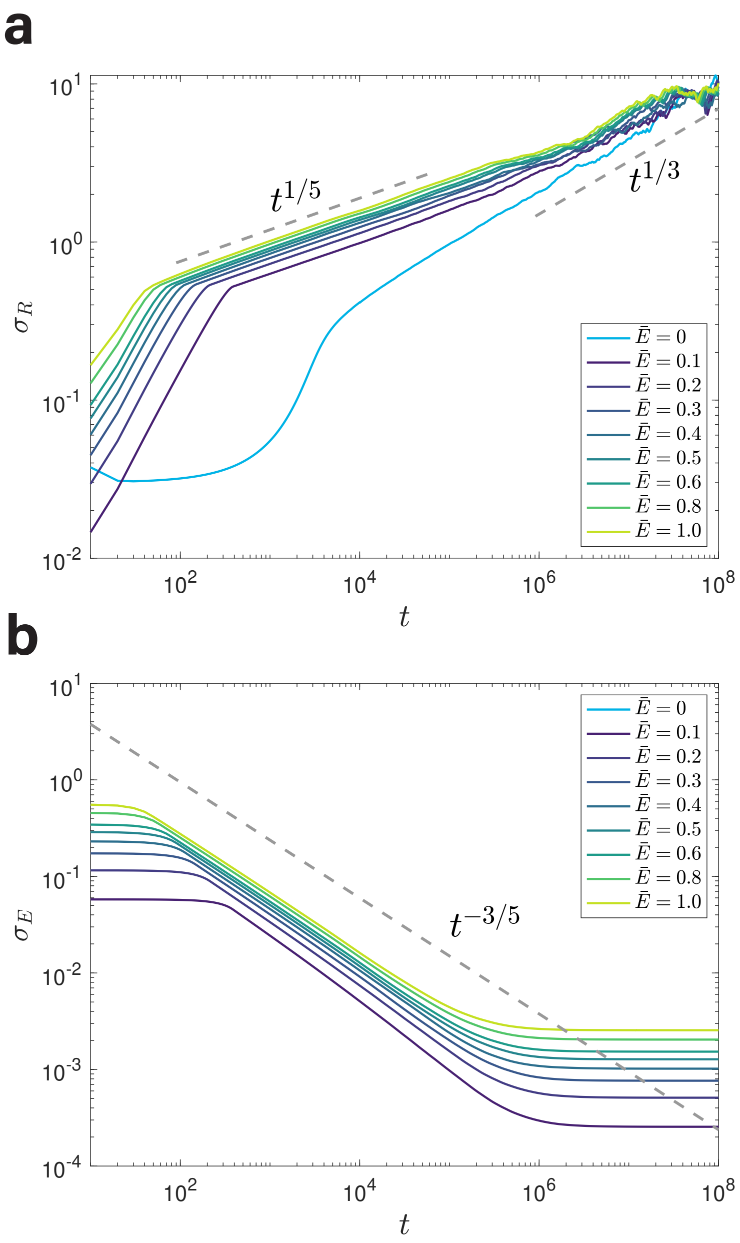


FIG. S5. Power-law scaling of $\sigma_R$ and $\sigma_E$. (a) $\sigma_R$ shows a power-law scaling with time, and the exponent is 1/5, which is different from the 1/3 scaling in Ostwald ripening as $\bar{E} = 0$. (b) $\sigma_E$ also shows a power-law scaling with time, and the exponent is $-3/5$.

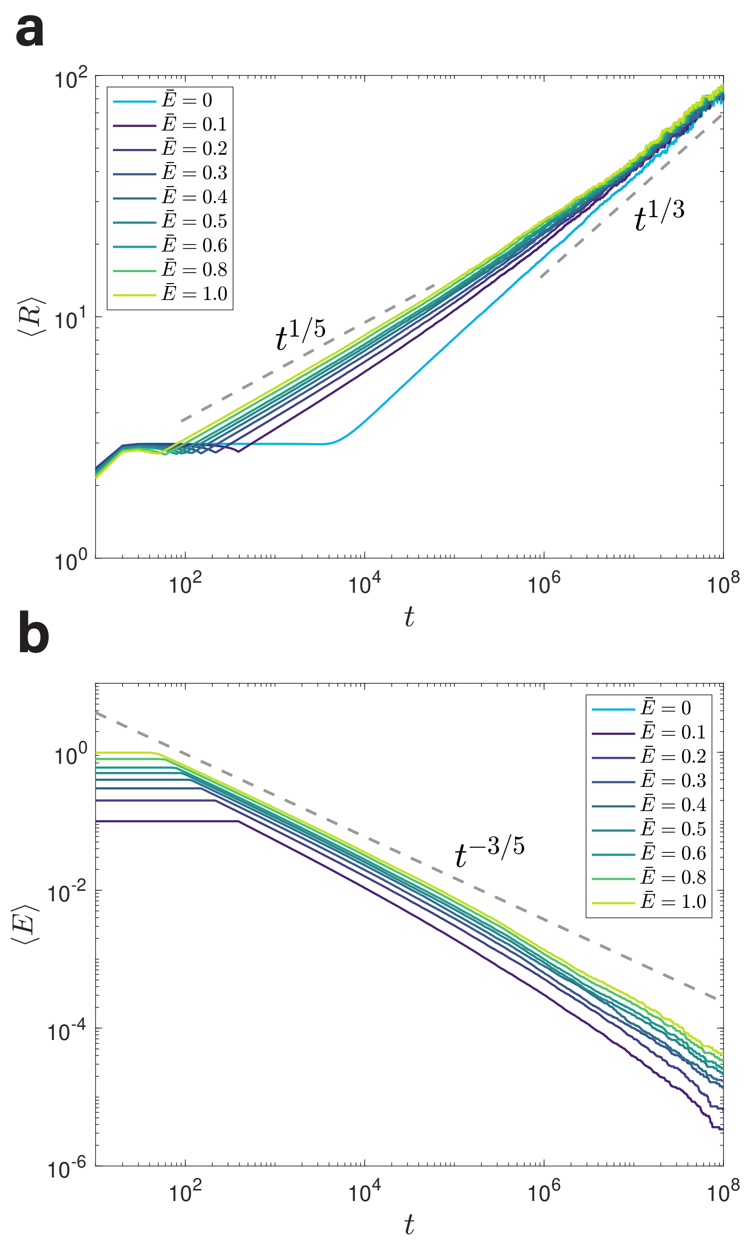


FIG. S6. Non-weighted average for $\langle R \rangle$ and $\langle E \rangle$. (a) The same analysis as Figure 2b in the main text where $\langle R \rangle$ is the non-weighted average by excluding dissolved condensates explicitly. (b) The same analysis as Figure 2c in the main text where $\langle E \rangle$ is the non-weighted average.

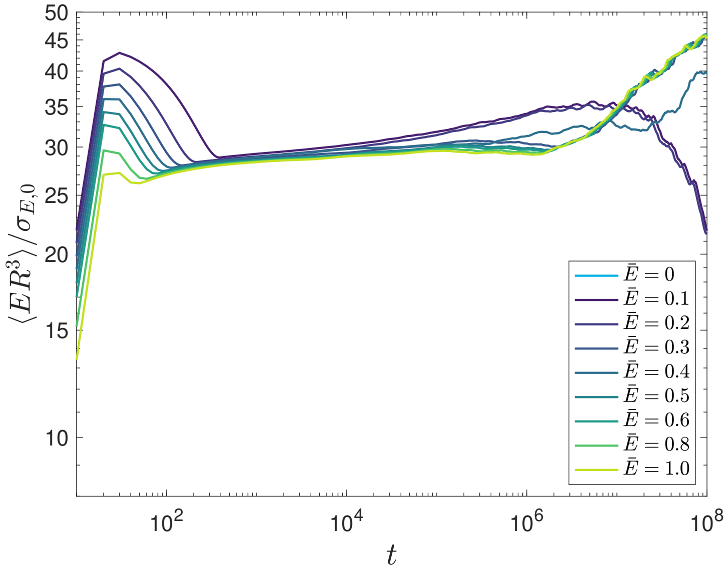


FIG. S7. The average elastic energy per condensate $\langle ER^3\rangle$ is nearly constant during the elastic ripening, meaning the elastic ripening process is scale invariant. Data of $\langle ER^3\rangle$ with different $\bar{E}$'s are collapsed after dividing the factor $\sigma_{E,0}$, indicating that $\sigma_{E,0}$ is the only scale of elastic energy.

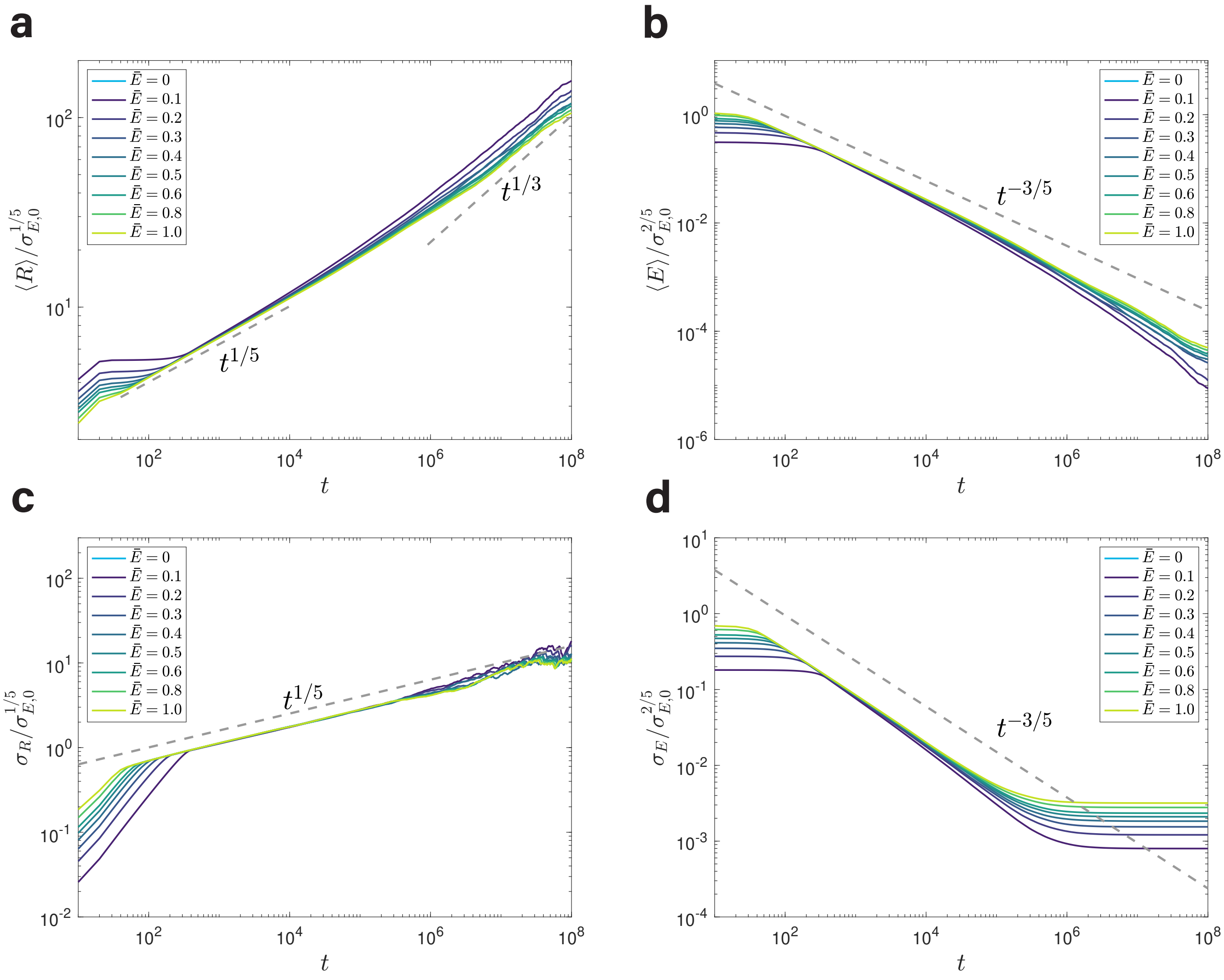


FIG. S8. Confirmation of the factors $A$ and $B$ in Eqs. (7, 8) of the main text. (a) $\langle R \rangle$ shows a power-law scaling with time, and the exponent is $1/5$. Results of different $\bar{E}$'s can be collapsed by dividing the factor $\sigma_{E,0}^{1/5}$. (b) $\langle E \rangle$ also shows a power-law scaling with time, and the exponent is $-3/5$. Results of different $\bar{E}$'s can be collapsed by dividing the factor $\sigma_{E,0}^{2/5}$. (c) The same analysis as (a) but for $\sigma_R$. (d) The same analysis as (b) but for $\sigma_E$.

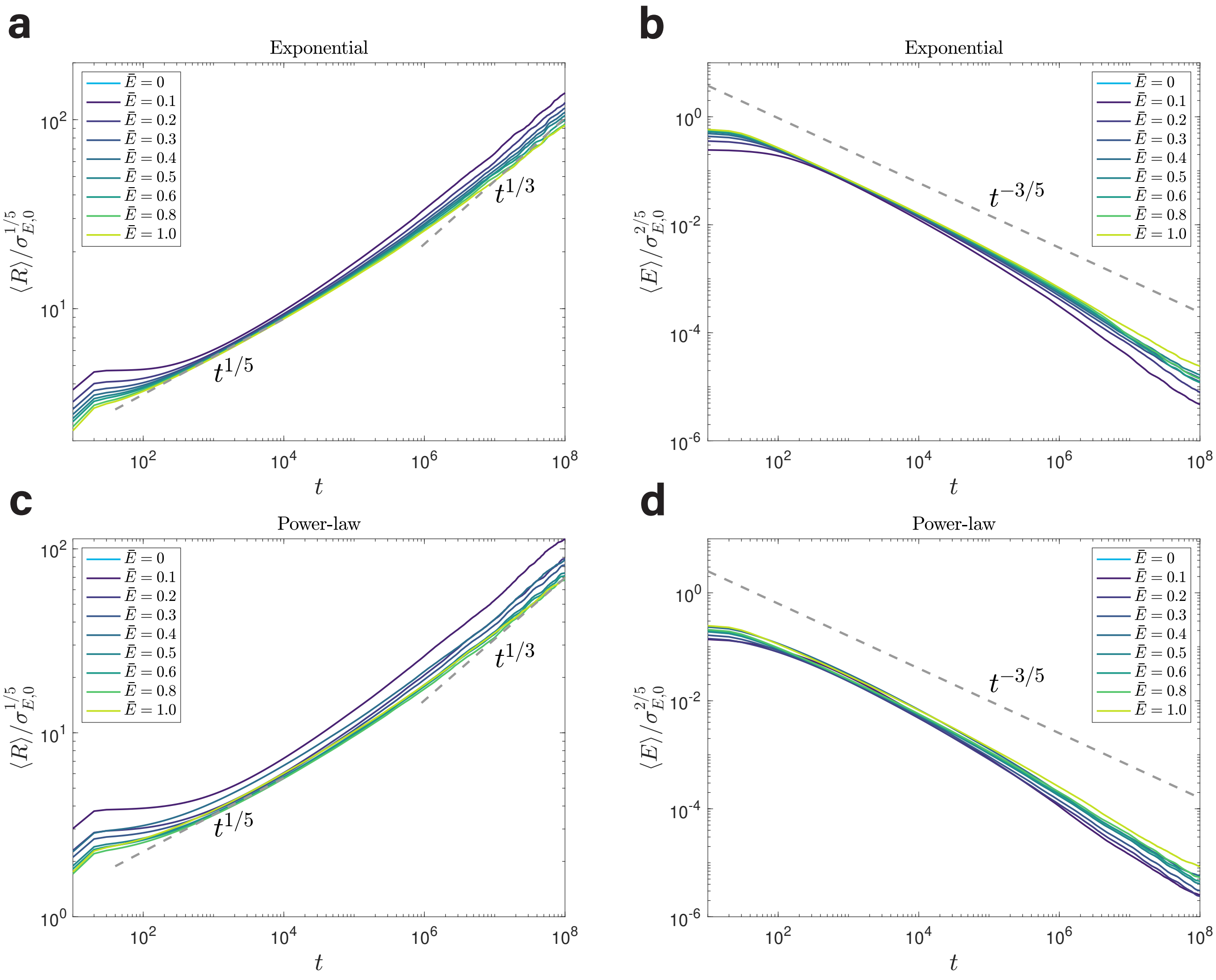


FIG. S9. Power-law scaling of $\langle R\rangle$ and $\langle E\rangle$ for different distributions of local elastic pressures. (a) In this case, $E$ follows an exponential distribution, $P(E) = \exp(-E/\bar{E})/\bar{E}$. The average radius $\langle R\rangle$ exhibits the predicted 1/5 scaling with the factor $\sigma_{E,0}^{1/5}$. (b) The average local pressure weighted by condensates' volumes also exhibits the predicted power-law scaling with the factor $\sigma_{E,0}^{2/5}$. (c) A similar analysis as (a) but with $E$ obeying a power-law distribution, $P(E) = 2((E+\bar{E})/\bar{E})^{-3}/\bar{E}$ with the average equals to $\bar{E}$. (d) The same analysis as (b) but for the power-law distributed $E$. In all panels, the number of nucleation sites $N = 5\times10^5$.

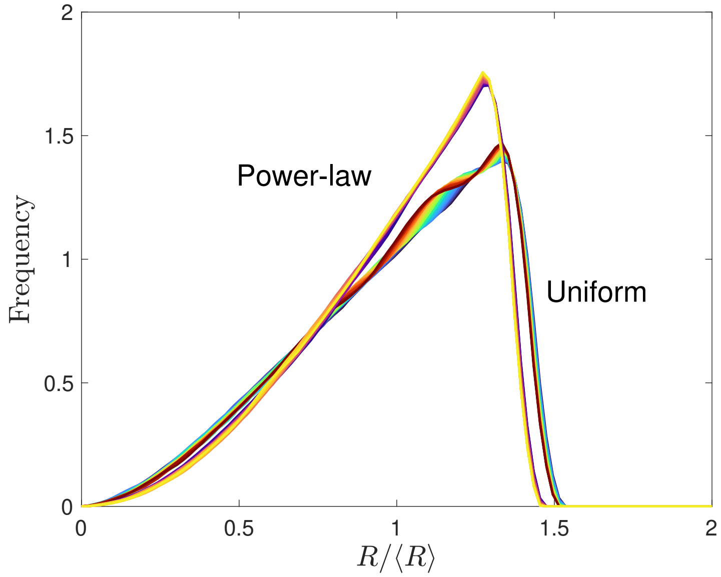


FIG. S10. Distributions of $R/\langle R\rangle$ for different distributions of $E$, including uniform and power-law distributions. In both cases, the distributions of $R/\langle R\rangle$ are self-similar between different times, although the shapes differ. The time is from $t = 4000$ to $t = 10000$ for both cases. In this figure, $\bar{E} = 1$, $R_n = 1.5R_c$. The number of nucleation sites $N = 5 \times 10^5$.

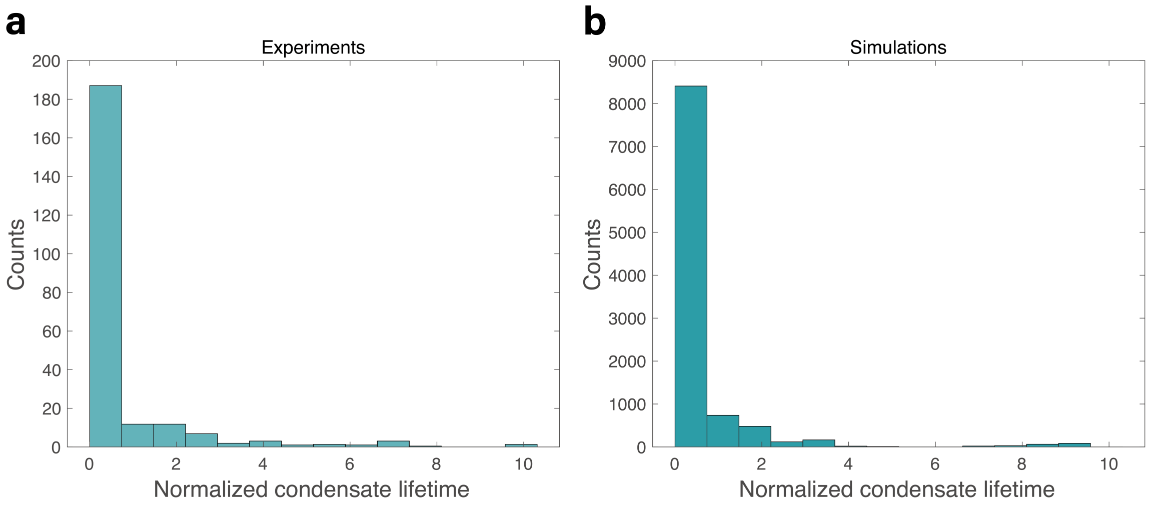


FIG. S11. Comparison between (a) experiments and (b) simulations of condensate lifetime distributions. The experimental data are from Ref. [6], which are the lifetime distribution of transient Pol II condensates in live mouse embryonic stem cells. Data of all the condensates that can initiate growth are used in the simulations. In both panels, the data are normalized with their averages over all condensates. In (b), $\bar{E} = 1$, the radii of nucleation sites $R_n = 5R_c$, $k_{\mathrm{dis}} = 10^{-6}$ and the number of nucleation sites $N = 10^5$.

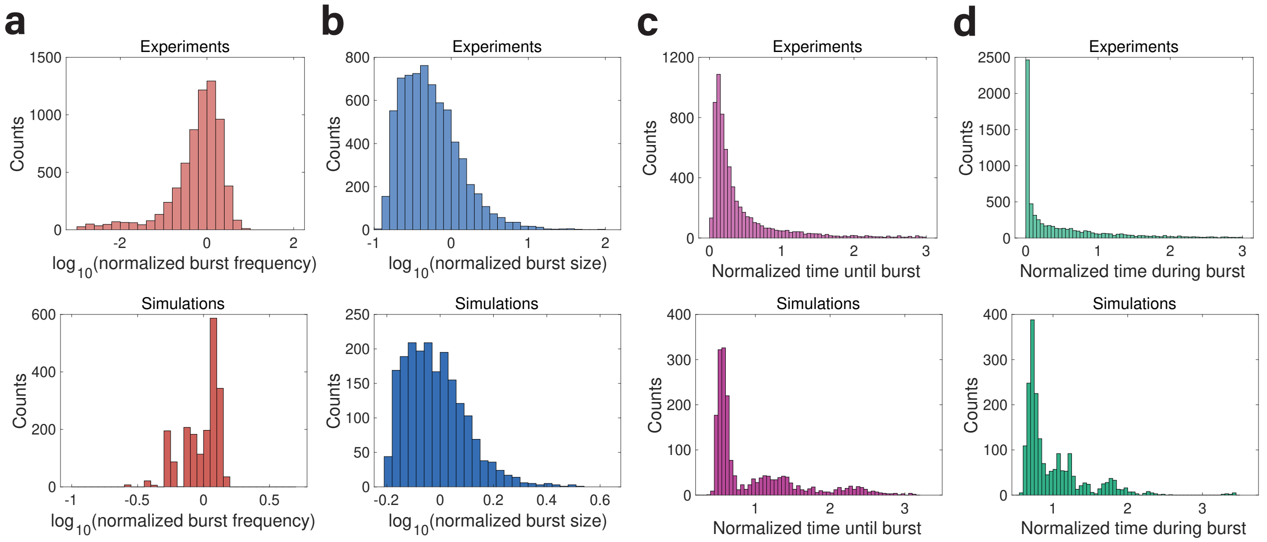


FIG. S12. The same analysis as Figure 5 in the main text, but with different simulation parameters. In all panels of simulations, $\bar{E} = 1$, the radii of nucleation sites $R_n = 2R_c$, $k_{\mathrm{dis}} = 10^{-5}$ and the number of nucleation sites $N = 10^5$. We exclude the data of condensates whose burst size is smaller than a threshold $10^5$.

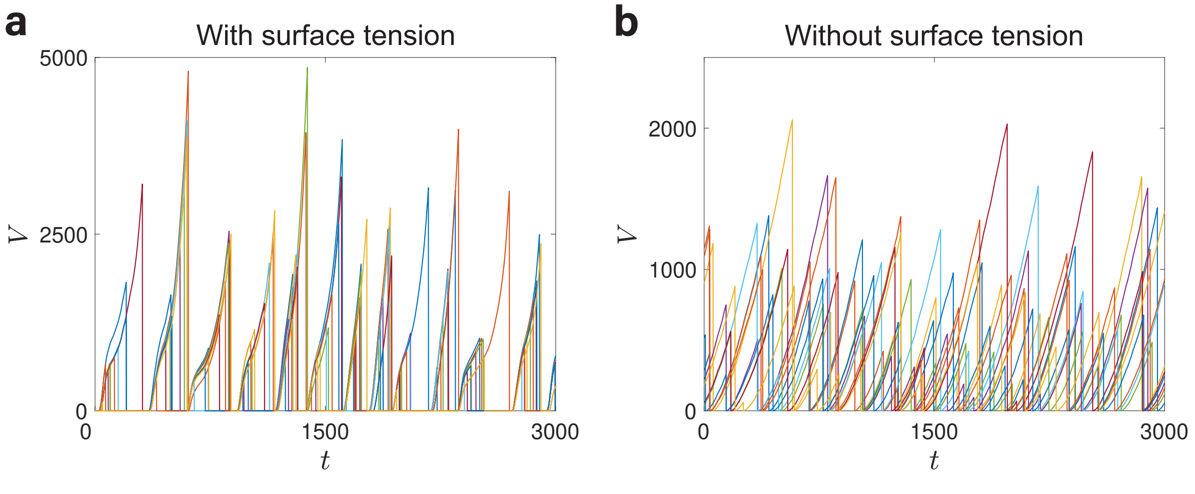


FIG. S13. Synchronization does not occur in the system without surface tension. (a) The same figure as Figure 6a in the main text, where all condensates are fully synchronized. (b) The same as (a) except that the surface tension does not contribute to the confining pressure. In this case, the growth of different condensates becomes completely independent, which means that synchronization is generated by surface tension. In both panels, $R_n = 1.2R_c$, $k_{\mathrm{dis}} = 10^{-5}$ and $\bar{E} = 1$. The number of nucleation sites $N = 2000$.

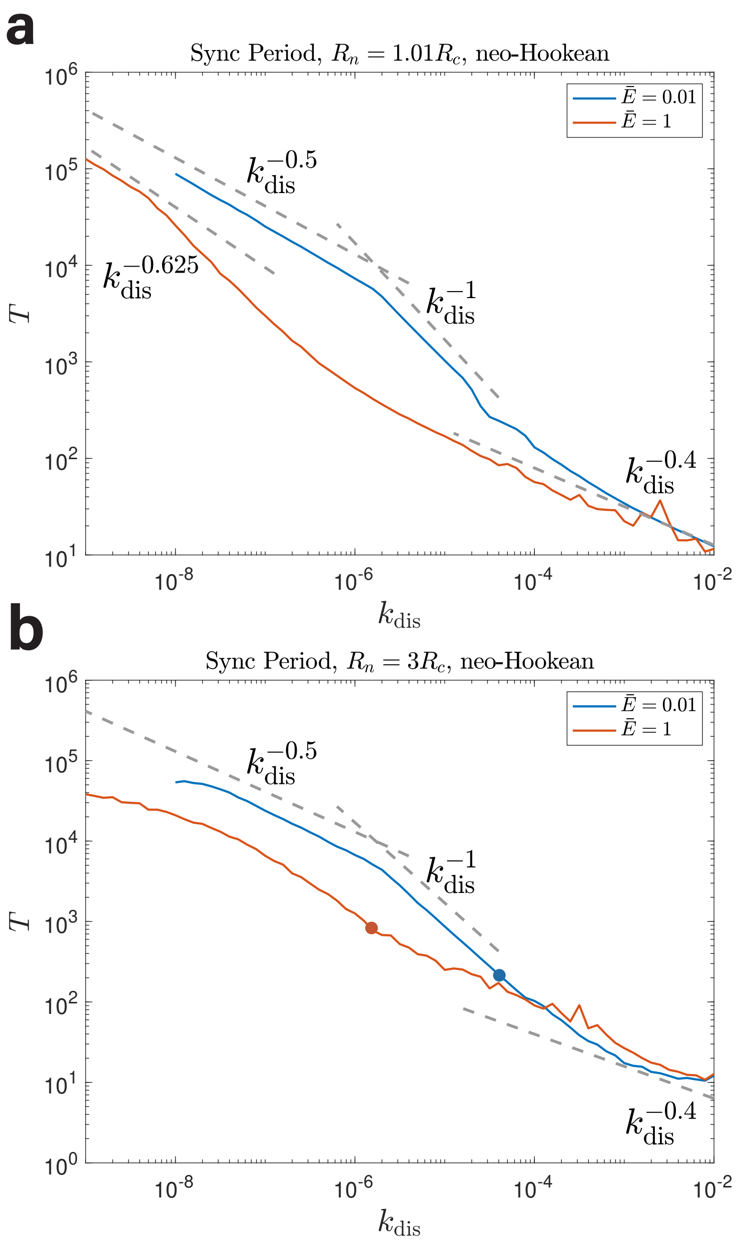


FIG. S14. The synchronization period $T$ as a function of the dissolution rate $k_{\rm dis}$. (a) In this panel, $R_n = 1.01R_c$ so that the synchronization fraction $f_{\rm sync} = 1$ in the entire range of $k_{\rm dis}$. In the case of $\bar{E} = 1$, some nucleation sites with large elastic pressure never grow during the simulation, so the volumes of growing condensates are larger than in the case of $\bar{E} = 0.01$. Since the dissolution rate per condensate is proportional to the volume, the period becomes smaller in the case of $\bar{E} = 1$. In the case of $\bar{E} = 1$, we see a signature of elastic ripening in which $T \sim k_{\rm dis}^{-0.625}$. (b) In this panel, $R_n = 3R_c$ so that the synchronization fraction $f_{\rm sync}$ can be smaller than 1 in the simulated range of $k_{\rm dis}$, which we use two circles to label the threshold $k_{\rm dis}$ at which $f_{\rm sync} = 1$. Note that the scaling relations $T \sim k_{\rm dis}^{-0.5}$ (or $T \sim k_{\rm dis}^{-0.625}$) and $T \sim k_{\rm dis}^{-1}$ only apply to a closed system of synchronized condensates, i.e., $f_{\rm sync} = 1$. Therefore, for $k_{\rm dis}$ above the threshold $k_{\rm dis}$, only the $T \sim k_{\rm dis}^{-0.4}$ scaling can be seen. In this figure, the number of nucleation sites $N = 2000$.

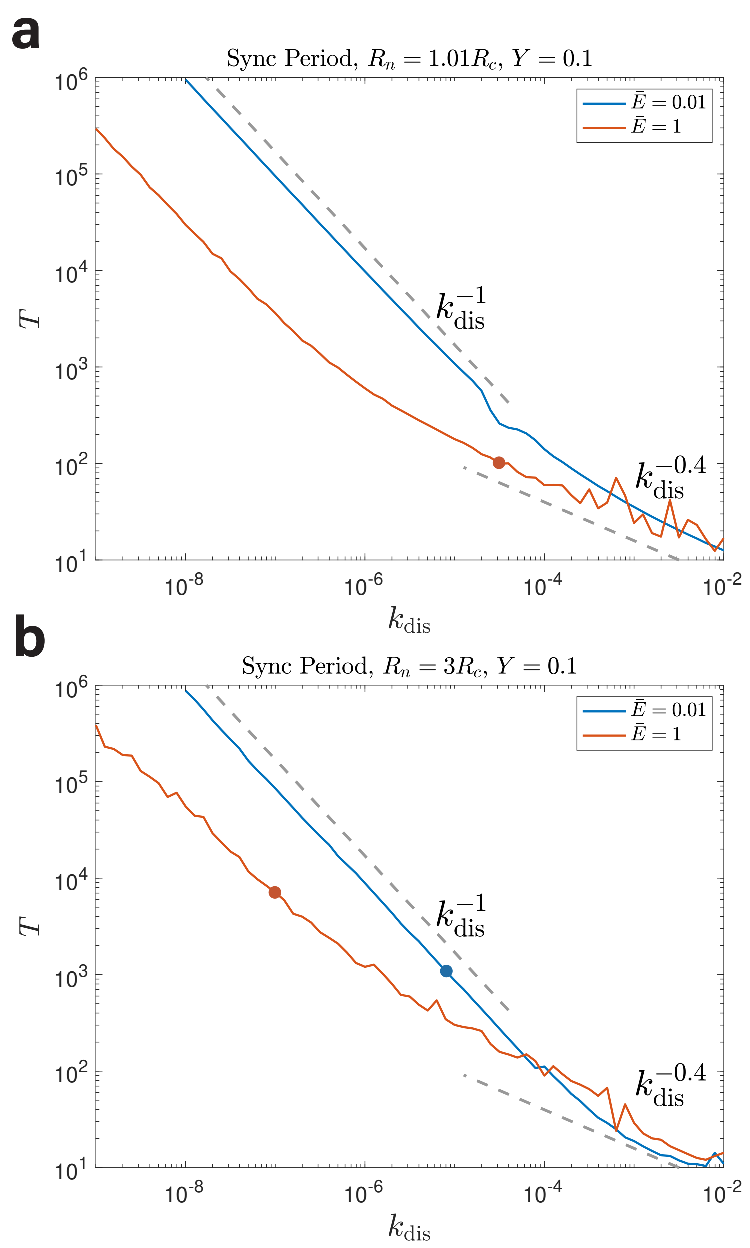


FIG. S15. The synchronization period $T$ as a function of the dissolution rate $k_{\mathrm{dis}}$ in an elastic medium beyond neo-Hookean elasticity. (a) In this panel, $R_n = 1.01R_c$. In the case of $\bar{E} = 1$, the fraction of synchronized condensates $f_{\mathrm{sync}}$ becomes smaller than 1 when $k_{\mathrm{dis}}$ is larger than the threshold value labeled by a circle. (b) In this panel, $R_n = 3R_c$ and we use two circles to label the threshold $k_{\mathrm{dis}}$ at which $f_{\mathrm{sync}} = 1$. Note that the scaling relation $T \sim k_{\mathrm{dis}}^{-0.5}$ (or $T \sim k_{\mathrm{dis}}^{-0.625}$) disappears in this model since the ripening process is suppressed in a medium beyond neo-Hookean. Meanwhile, because the condensate volume will reach a plateau value simply due to the increasing elastic pressure, the $T \sim k_{\mathrm{dis}}^{-1}$ scaling does not require $f_{\mathrm{sync}} = 1$ any more. In this figure, the number of nucleation sites $N = 2000$.
